# Vegetation structure determines cyanobacterial communities during soil development across global biomes

**DOI:** 10.1101/2021.09.11.459883

**Authors:** Concha Cano-Díaz, Fernando T. Maestre, Juntao Wang, Jing Li, Brajesh Singh, Victoria Ochoa, Beatriz Gozalo, Manuel Delgado-Baquerizo

## Abstract

- Soil cyanobacteria play essential ecological roles and are known to experience large changes in their diversity and abundance throughout early succession. However, much less is known about how and why soil cyanobacterial communities change as soil develops from centuries to millennia, and the effects of aboveground vegetation on these communities.
- We combined an extensive field survey including 16 global soil chronosequences across contrasting ecosystems (from deserts to tropical forests) with molecular analyses to investigate how the diversity and abundance of soil cyanobacteria under vegetation change during soil development from hundreds to thousands of years.
- We show that, in most chronosequences, the abundance, species richness and community composition of soil cyanobacteria were relatively stable as soil develops (from centuries to millennia). Regardless of soil age, forest chronosequences were consistently dominated by non-photosynthetic cyanobacteria (Vampirovibrionia), while grasslands and shrublands were dominated by photosynthetic cyanobacteria. Chronosequences undergoing drastic vegetation shifts during soil development (e.g. transitions from grasslands to forests) experienced significant changes in the composition of soil cyanobacteria communities.
- Our results advance our understanding of the ecology of cyanobacterial classes, specially the understudied non-photosynthetic ones and highlight the key role of vegetation as a major driver of their temporal dynamics as soil develops.

## Introduction

Cyanobacteria are a diverse and functionally important group of microorganisms on Earth. This phylum includes microbes with a wide range of metabolic capabilities from the photosynthetic cyanobacteria (class Cyanobacteriia), which live in most types of illuminated environments (Whitton & Potts, 2012), to non-photosynthetic clades such as Vampirovibrionia and Sericytochromatia (Shih *et al*., 2017; Soo *et al*., 2017). A wide variety of life strategies are included in the phylum, from autotrophic to parasitic ones including multiple symbioses with plants and animals (Lindo *et al*., 2013; Soo *et al*., 2015, Benavent-Gonzalez 2019). In soil environments, cyanobacteria are globally distributed (Moreira *et al*., 2013; Cano-Díaz *et al*., 2020) and have an estimated biomass that can reach 54 × 10^12^ gC (Garcia-Pichel *et al*., 2003). They are also important components of biological soil crusts (Büdel *et al*., 2016), which cover up to 12% of Earth’s terrestrial surface (Rodriguez-Caballero *et al*., 2018).

Soil cyanobacterial communities are known to undergo profound changes in their abundance, diversity and composition during the first days to decades of primary and secondary successions(Kastovská *et al*., 2005; Nemergut *et al*., 2007; Schmidt *et al*., 2008a; Dojani *et al*., 2011; Arróniz-Crespo *et al*., 2014). Filamentous non-heterocystous (without heterocytes) cyanobacteria as *Microcoleus vaginatus* appear first and contribute to soil stabilization through their rope-like bundles and exopolysaccharide sheath that binds soil particles together (Mazor *et al*., 1996; Garcia-Pichel & Wojciechowski, 2009). Then, heterocystous nitrogen-fixers become more abundant and contribute to enhance soil microhabitat conditions, facilitating the colonization and further development of vegetation and other organisms (Danin *et al*., 1998; Garcia-Pichel & Wojciechowski, 2009; Singh *et al*., 2016). When this happens, the abundance of cyanobacteria decreases and they no longer constitute the dominant fraction of the soil (Schmidt *et al*., 2008b; Maier *et al*., 2018). Despite the fundamental ecological roles they play, much less is known about how cyanobacterial communities change as soil develops from centuries to millennia, and their relationship with vegetation structure changes with time. Furthermore, non-photosynthetic classes are particularly understudied (Soo *et al*., 2014, 2017) and because of their inferred metabolic capabilities, their abundance could increase with soil development due to the raise of organic matter or by the opening of new ecological niches (Laliberté, 2014; Laliberté *et al*., 2017), but empirical evidence of this is largely lacking.

Direct observation and experimental manipulations in natural ecosystems over long time periods are not easy to obtain, therefore the use of long-term chronosequences is a suitable indirect alternative to explore them (Walker *et al*., 2010; Blois *et al*., 2013). Soil chronosequences imply a set of sites from the same parental material forming a spatial gradient that represents different stages of soil development. These space-for-time substitutions, which have successfully been used to characterize temporal changes in multiple ecological processes (Bardgett *et al*., 2005; Blois *et al*., 2013), have demonstrated that vegetation structure and soil nutrient contents shift during ecosystem development. For example, plant cover, soil carbon and nitrogen, which are mainly associated to biological activity, are known to increase as soil develops (Vitousek, 2004; Wardle *et al*., 2004; Laliberté, 2014), while soil phosphorous is known to decrease over millennia due to weathering and leaching (Vitousek, 2004). The main groups of photosynthetic and non-photosynthetic cyanobacterial taxa are known to be strongly associated with vegetation structure, soil pH and nutrient content at the global scale (Dini-Andreote *et al*., 2014; Cano-Díaz *et al*., 2020), and therefore, might also undergo through predictable changes in diversity, abundance and composition as soil develops from hundreds to thousands of years. However, this has not been tested in the field yet.

Here, we investigate how the abundance, diversity and composition of soil cyanobacterial communities under vegetation change as soil develops from hundreds to thousands of years across contrasting ecosystem types. We used the most comprehensive soil chronosequence standardized survey conducted to date (Delgado-Baquerizo *et al*., 2019), which includes 16 global chronosequences from polar ecosystems and deserts to temperate and tropical forests and encompass a wide range of climatic conditions, vegetation types and soil ages (from 10 to 5*10^6^ years). This database has been previously used to identify the overall changes in the richness (number of species) of soil bacteria, fungi, protists and invertebrates (Delgado-Baquerizo *et al*., 2019). However, the changes in the diversity and abundance of soil cyanobacteria remains unexplored. Using this unique chronosequence survey, we tested the following hypotheses:

### H1. Soil age indirectly regulates soil cyanobacterial communities

We hypothesized that relatively young and sparsely vegetated soils (from years to decades) could be potentially dominated by Cyanobacteriia and specifically by nitrogen-fixing cyanobacteria. We expect this because, unlike for nutrients associated with bedrock (e.g., phosphorus), young soils are limited by organic carbon and nitrogen (Hooper *et al*., 2000; Wardle *et al*., 2004), which are fixed by biological communities (Belnap, 2002; Hayat *et al*., 2010). This way, diazotroph-dominated cyanobacterial communities might play critical roles in the functioning of relatively young ecosystems (Brankatschk *et al*., 2011). As soils develop, with the accumulation of nutrients and the availability of new potential niches linked to vegetation succession (Van der Putten *et al*., 2013), we expect to find an increase in the abundance of non-fixing N cyanobacteria and the switch to more heterotrophic communities including non-photosynthetic cyanobacteria (Laliberté *et al*., 2017).

### H2. Biome type determines the structure of soil cyanobacterial communities regardless of soil age

Climate and ecosystem-level soil properties and vegetation might uniquely influence the abundance, composition and diversity of soil cyanobacteria regardless of soil age. For example, we would expect low light conditions and acid soils from tropical and temperate forests to harbor different cyanobacterial communities than those from deserts or cold ecosystems. There, adaptations to dessication and high sun exposure due to the sparse or minimum vegetation are expected. Similarly, climatic conditions, rather than soil age, are expected to regulate the proportion of cyanobacterial with different functional capabilities e.g, non-photosynthetic vs, photosynthetic cyanobacteria (Cano-Díaz *et al*., 2020).

## Materials and Methods

### Chronosequence survey and soil analyses

We obtained cyanobacterial 16S rRNA gene sequences from a global survey of 16 soil chronosequences from six continents conducted between 2016 and 2017 (Delgado-Baquerizo *et al*., 2019) (Fig.1). These chronosequences include a wide range of soil ages (from centuries to millennia), vegetation types (grasslands, shrublands, forests and croplands), origins (volcanic, sedimentary, dunes and glacier) and climatic types (tropical, temperate, continental, arid and polar).

**Figure 1.**
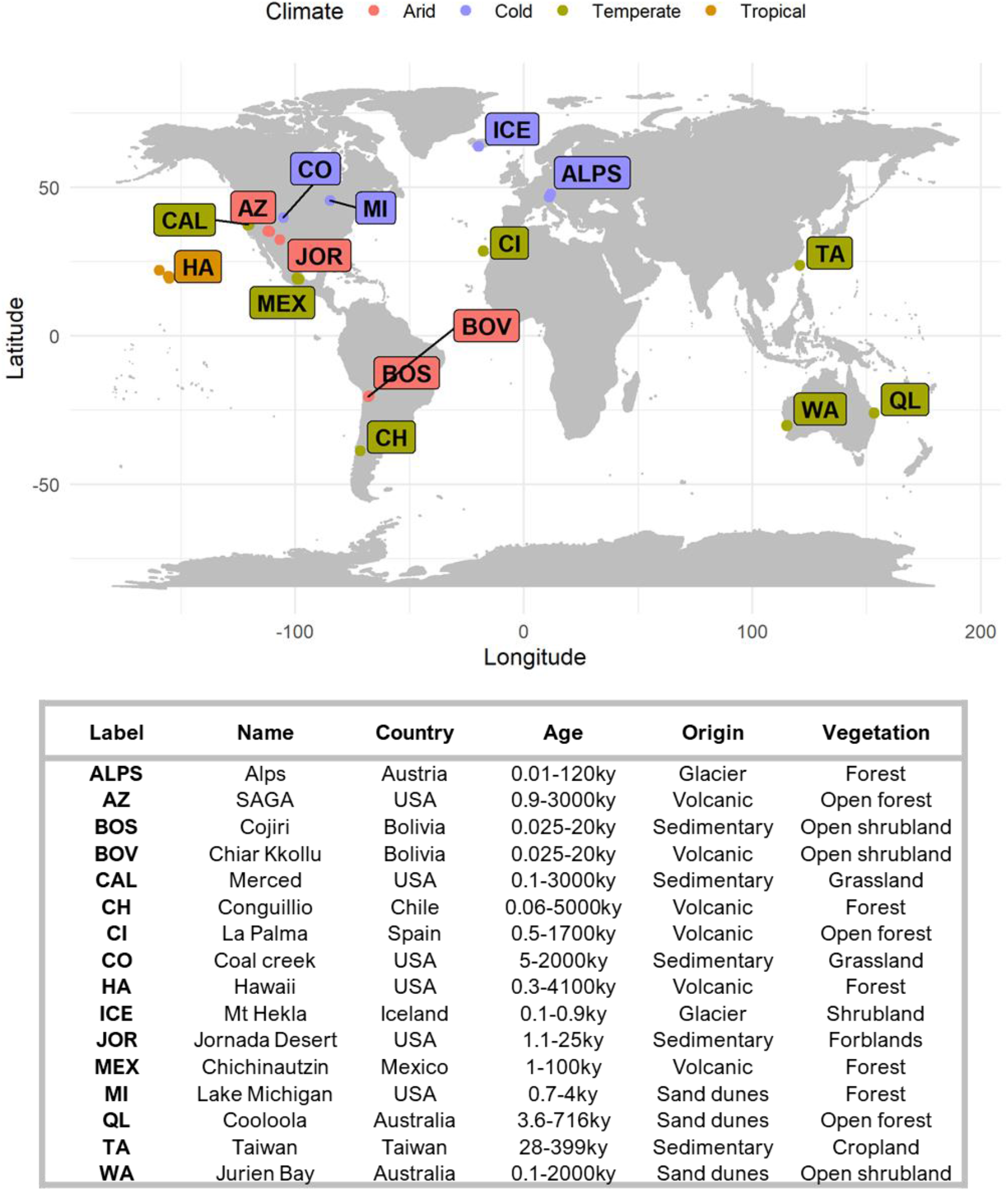
Locations of the chronosequences used in this study.

All field surveys were conducted following a standardized sampling protocol, with a plot size selected to accommodate the different plant sizes and ecosystem types covered in our study. Each chronosequence was divided in 4-10 stages related to soil age (soil age was determined with radiometric dating measurements, references and measurements in Supplementary Tables ST1 and ST2; further details in Delgado-Baquerizo *et al*., 2019). Each stage was sampled using a 50 × 50 m plot with three parallel transects of 50 m spaced apart 25 m. Total plant cover and the number of perennial plant species (plant richness) was measured in each transect with the line-point intercept method (Tongway & Hindley, 2004). We obtained the standardized average of mean temperature (MAT) and precipitation (MAP) per plot using climatic data from www.worldclim.org. Aridity Index (precipitation/potential evapotranspiration) was obtained from the Global Potential Evapotranspiration database (Zomer *et al*., 2008), which uses interpolations from WorldClim. At each plot, five composite soil samples (0-10 cm depth) were collected under the dominant vegetation and sieved (2 mm) back in the laboratory. The collection depth could be diluting the abundance of light dependant photosynthetic cyanobacteria in favor to the non-photosynthetic classes. However since soils are always collected at the same depth, relative increases and decreases in cyanobacterial types with time are directly comparable. A fraction of the sieved soil was immediately frozen at −20°C for molecular analyses; the other fraction was air-dried for biochemical analyses. We retrieved 10 g from each frozen composite soil sample and ground it with a mortar and liquid N to obtain a homogenized representative sample. Soil DNA was extracted afterwards with the PowerSoil DNA Isolation Kit (MoBio Laboratories, Carlsbad, CA, USA), following manufacturer’s instructions.

All soil analyses were done in all samples in Rey Juan Carlos University (Spain) to facilitate the comparison of results. Soil pH was measured in a 1:2:5 suspensions of dry mass to deionized water volume with a pH meter, and ranged from 3.19 to 9.45. The concentration of soil total organic C (soil C hereafter) was determined by colorimetry after oxidation with a mixture of potassium dichromate and sulfuric acid (Anderson & Ingram, 1989) and ranged from 0.3 to 473.6 g CKg-1. Available P was determined from bicarbonate extracts as described in (Olsen & Sommers, 1982) and ranged from < 0.01 to 90.69 mg P kg^-1^. Total N was measured with a CN analyzer (LECO CHN628 Series LECO Corporation, St Joseph, MI, USA) and ranged from <0.01 to 1.379%. Further details are described in Delgado-Baquerizo *et al*. (2019)

### Cyanobacterial abundance, richness and composition

To obtain the total abundance of cyanobacteria, quantitative PCR (qPCR hereafter) was performed with specific cyanobacterial primers CYA 359F and CYA 781 R(a) and CYA 781 R(b) (Nübel *et al*., 1997). Each qPCR reaction contained 5 ul of LightCycler^®^ 480 SYBR Green I Master, 0.5 ul of forward primer CYA 359, 0.25 ul of CYA 781 R(a) and 0.25 ul of CYA 781 R(b) reverse primers, 1 ul of template and 3 ul of sterilized water. Reaction conditions included an initial denaturalization stage of 95°C for 5 minutes, followed by 40 cycles of amplification: 95°C for 1 min, 50°C for 1 min and 72°C for 1 min; and finally a melting curve of 95°C for 5s and 65°C for 1 min. The qPCR machine used was a Roche light cycler 480 (Roche Diagnostics, Mannheim, Germany), that uses the Crossing point value, where your sample crosses the threshold fluorescence so you can infer its abundance with standard samples. Cyanobacterial abundance is expressed as 16S copies/g soil. All qPCR analyses were done at the facilities of Western Sydney University (Australia).

The primers 515F-806R (Caporaso *et al*., 2010b) were used to amplify the bacterial V4 region of the 16S SSU rRNA using the Illumina Miseq platform at the Next Generation Genome Sequencing Facility at Western Sydney University (Australia). Bioinformatic processing was performed using a combination of QIIME (Caporaso *et al*., 2010a), USEARCH (Edgar, 2010) and UNOISE2 (Edgar, 2016). Phylotypes (i.e Operational Taxonomic Units; OTUs) were identified at the 100% identity level (zOTUs). Taxonomy was assigned with the online tool Silva Incremental Alligner with Silva database (Pruesse *et al*., 2012) according to last update (SILVA SSU 138.1), data accessed on 10 March of 2021. The 16S zOTU abundance table was rarefied and the relative abundance (%) of cyanobacteria in relation to total bacterial (16S rRNA) reads was calculated for community analyses. For diversity estimations, Shannon Index (Shannon & Weaver, 1949) was calculated using the *estimate_richness* function from the phyloseq R package v1.16.2 (Mcmurdie & Holmes, 2012). Phylotype richness was highly correlated with the Shannon Index (Spearman ρ =0.91, p<0.05), so we used it as a surrogate of phylotype diversity. Relative abundances (%) of cyanobacterial classes and families were calculated for statistical analyses; and sqrt-transformed to improve their goodness of fit to the normal distribution.

### Statistical analyses

We tested the relationship between cyanobacterial abundance and soil development (chronosequence stage and soil age) with linear mixed-effects models. Between both, the stage of the chronosequence was selected because models have larger percentage of variance explained (R^2^) than those with soil age (the radiometric dating estimation in years); although similar results were obtained with both approximations.

Linear mixed-effects models were used to analyze the total abundance of cyanobacteria, phylotype richness, and the relative abundances of cyanobacterial classes with stage of the chronosequence as a fixed factor. These analyses allowed us to obtain generalizable results as they consider the chronosequence as a random factor, thus controlling dependencies between plots within a chronosequence. Models were done with *lmer* function from the lme4 R package (Bates *et al*., 2015) and tests for significance were calculated with lmertest R package (Kuznetsova *et al*., 2017). To further explore the relationships between abundance and phylotype richness through time, linear and polynomic (quadratic and cubic) regression models were fitted. These analyses were performed with the *lm* function from base R keeping only relationships with P < 0.05 and selecting the model with the lowest Akaike Information Criterion (AIC) value (Akaike, 1974).

We analyzed the changes in the composition of soil cyanobacteria between stages separately for each of the 16 chronosequences surveyed using multivariate PERMANOVAs (Anderson, 2001). These analyses were conducted with Unifrac-weighted distances and the phyloseq R package (Mcmurdie & Holmes, 2012). Unifrac-weighted distances are more informative than non-phylogenetic distance measures because they take into consideration the degree of similarity between phylotype-sequences and the relative abundance of each phylotype within samples (*weight* parameter). In those chronosequences with community changes, subsequent PERMANOVA analyses were done to evaluate the effect of vegetation (through total and tree covers). To determine whether the relative abundances of all cyanobacterial classes and families change between stages, we used multiple univariate PERMANOVAs with Euclidean distances. All PERMANOVA analyses were made with the *adonis* function and 1000 permutations of the vegan package (Oksanen, 2015)

We investigated the associations between cyanobacterial composition and environmental variables through statistical modelling (SEM) and ordination methods (PCoA). Principal Coordinate Analysis (PCoA) with Unifrac distances was computed to compare the composition of microbial communities among sites using the phyloseq package (Mcmurdie & Holmes, 2012). To show the influence of environmental variables in the composition of the cyanobacterial community, we made a selection of variables covering vegetation, climate and soil and fitted them into the ordination with *envfit* function and 9999 permutations with vegan package (Oksanen, 2015). We tested 11 variables including spatial dissimilarity (Euclidean distance between plots), soil age (years), vegetation (tree and grass cover and vascular plant richness), climate (MAT and MAP) and soil (pH, Soil C, Soil CN and Soil NP) properties. To be included in the analysis, these variables should not be strongly correlated among themselves (Pearson ρ<0.6 in all cases). To improve normality, we transformed response variables using either square root (abundance of cyanobacteria, relative abundance of the cyanobacterial classes and plant richness) or natural logarithm (soil age, Soil C, Soil CN and Soil NP) transformations. Since we are comparing variables that are in different scales, and to avoid over or underestimations in our model, all variables included were centred (by subtracting the variable means) and scaled (by dividing the variables by their standard deviations) with the *scale* function of the R Base package (Team, 2013). Because of the large number of variables tested, a Bonferroni correction test was used with the function *p*.*adjust* from stats package (Team, 2013) that resulted in the exclusion of MAT and Soil age from the analyses.

Structural equation modelling (SEM) was used to evaluate the direct and indirect effects of the different predictors of the abundance of cyanobacteria and of the relative abundance of the cyanobacterial classes. These include the same 11 variables used in the PCoA ordination and spatial dissimilarity (Euclidean distance between plots) was included to control for spatial autocorrelation of plots within the model. To test the overall fit of the model, we used Chi-square fit test and Root Mean Square Error of Approximation (RMSEA). The model has a good fit when Chi-square/df is low, (i.e. <2) and P is high (P>0.05) and RMSEA is indistinguishable from zero with a high P>0.05. All our indices suggested our model had an adequate model fit so we could interpret path coefficients. Each path coefficient is analogous to a regression weight and describes the strength and sign of the relationships between two variables. To deal with non-normally distributed variables, we used 5000 bootstraps to simultaneously test the significance of each path coefficient. Standardized total effects (direct plus indirect) of all predictors on each cyanobacterial abundance were calculated. SEM analyses were conducted with AMOS 24.0.0 (IBM SPSS, Chicago, IL, USA). The rest of statistical analyses were carried out using R version R-3.6.0 (Team, 2013).

## Results

### Changes in the abundance and diversity of soil cyanobacteria during soil development

Cyanobacterial abundance and phylotype richness ranged from 1.56 × 10^7^ to 1.83 × 10^9^ copies/g soil and from 1 to 50 phylotypes per sample, respectively. We found no consistent and generalizable effects of chronosequence stage on the richness and abundance of cyanobacteria across contrasting ecosystem types. In general, the richness and abundance of cyanobacteria showed very little and variable association with soil development (see Supplementary Table 3, Fig. S1), remaining stable through time across the contrasting ecosystem types studied. However, we found some associations between chronosequence stage and cyanobacterial diversity and abundance in three and four of the 16 chronosequences evaluated, respectively (Fig. 2). For example, we found an increase in the abundance of these organisms in relatively old temperate chronosequences from Mexico (adjusted R^2^= 0.15) and Australia (Jurien Bay, adjusted R^2^= 0.20). Nonlinear relationships between cyanobacterial abundance and soil age were found in Coal Creek and Merced chronosequences with the significant fitting of cubic functions (adjusted R^2^= 0.18 and 0.53 respectively) (Fig. 2a and Supplementary Table 4). Cyanobacterial richness decreased with time in the semiarid Substrate Age Gradient of Arizona (SAGA) chronosequence (adjusted R^2^=0.41). Nonlinear cubic relationships between richness and chronosequence stage were found in temperate chronosequences from Chile and Australia (Jurien Bay; adjusted R^2^= 0.41 and 0.53 respectively) (Fig. 2b and Supplementary Table 5). Jurien Bay was the only chronosequence experiencing changes in both abundance and diversity of cyanobacteria through time.

**Fig.2.**
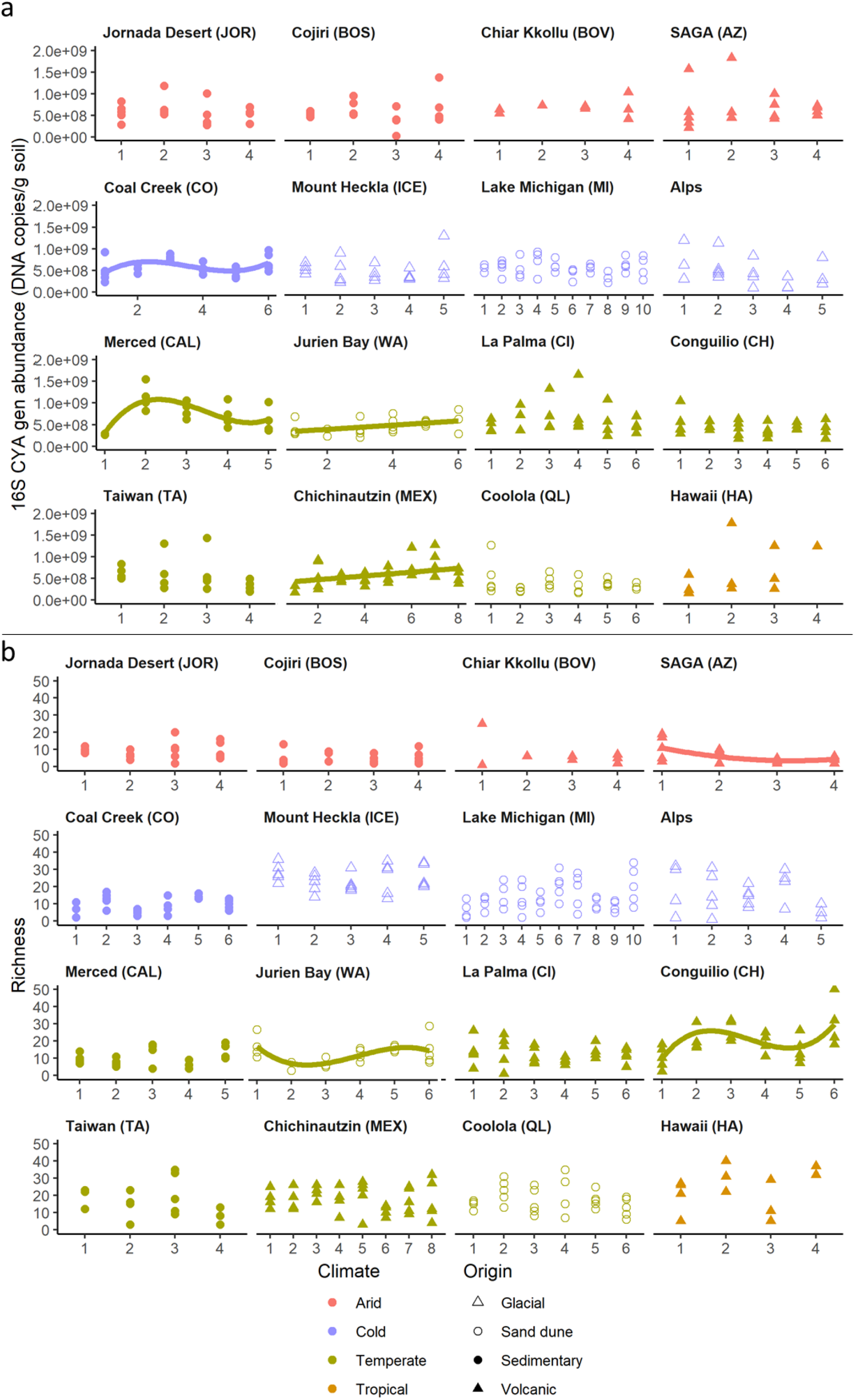
Cyanobacterial abundance (a) and richness (b) through soil development (chronosequence stage) for the 16 chronosequences studied. Lines show significant (p <0.05) model fits between chronosequence stage and cyanobacterial abundance (the fitted model is that with the lowest AIC value among linear, quadratic and cubic models). Details of all fitted models can be found in Supplementary Tables 3 and 4.

### Changes in the composition of soil cyanobacterial communities are associated with vegetation changes as soil ages

In general, the relative abundance of the Sericytochromatia class decreased with chronosequence stage, whereas the other cyanobacterial classes showed variable responses depending on the chronosequence considered (Fig S1, Supplementary Table 3). Most chronosequences were consistently dominated by one of the main ecological groups of cyanobacteria (Cyanobacteriia or Vampirovibrionia classes) over time (Fig. 3). However, changes in the cyanobacterial composition through time were found in Alps, Conguilio, Lake Michigan and Jurien Bay chronosequences (Fig. 3 and Supplementary Table 6). Differences in the composition of the cyanobacterial community between stages were noticeable even at a high taxonomic level-class level-in these chronosequences. This was confirmed with univariate PERMANOVA analyses in all cases except for Conguilio. A particular turnover pattern was observed in Jurien Bay, Alps and Lake Michigan, where early stages were dominated by photosynthetic cyanobacteria (Cyanobacteriia) and late ones by nonphotosynthetic Vampirovibrionia (Fig. 3). At the family-level, shifts through chronosequence stage in the relative abundances of cyanobacterial families were also observed in chronosequences with significant community changes (see Fig. 4). The relative abundance of N-fixers from the Nostocaceae family decreased as soil aged in Jurien Bay and Alps chronosequences. Similar responses were observed in the Nodosilineaceae family in Lake Michigan and Alps chronosequences.

**Fig.3.**
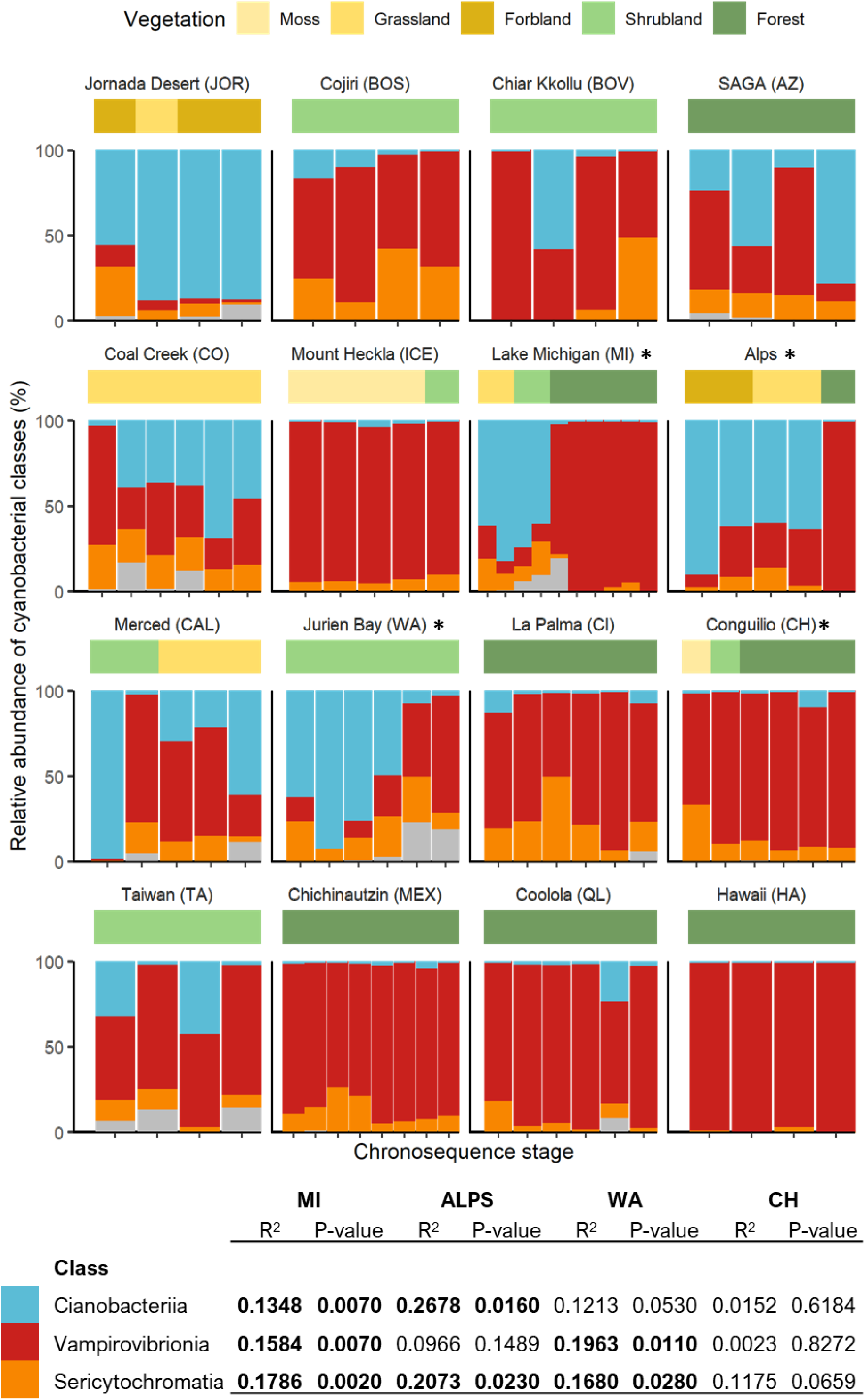
Relative abundance of cyanobacterial classes (%) and vegetation types across stages for all chronosequences. Bottom table: Summary of PERMANOVA results for relative abundances of cyanobacterial classes in chronosequences with large changes in cyanobacterial composition.

**Fig. 4:**
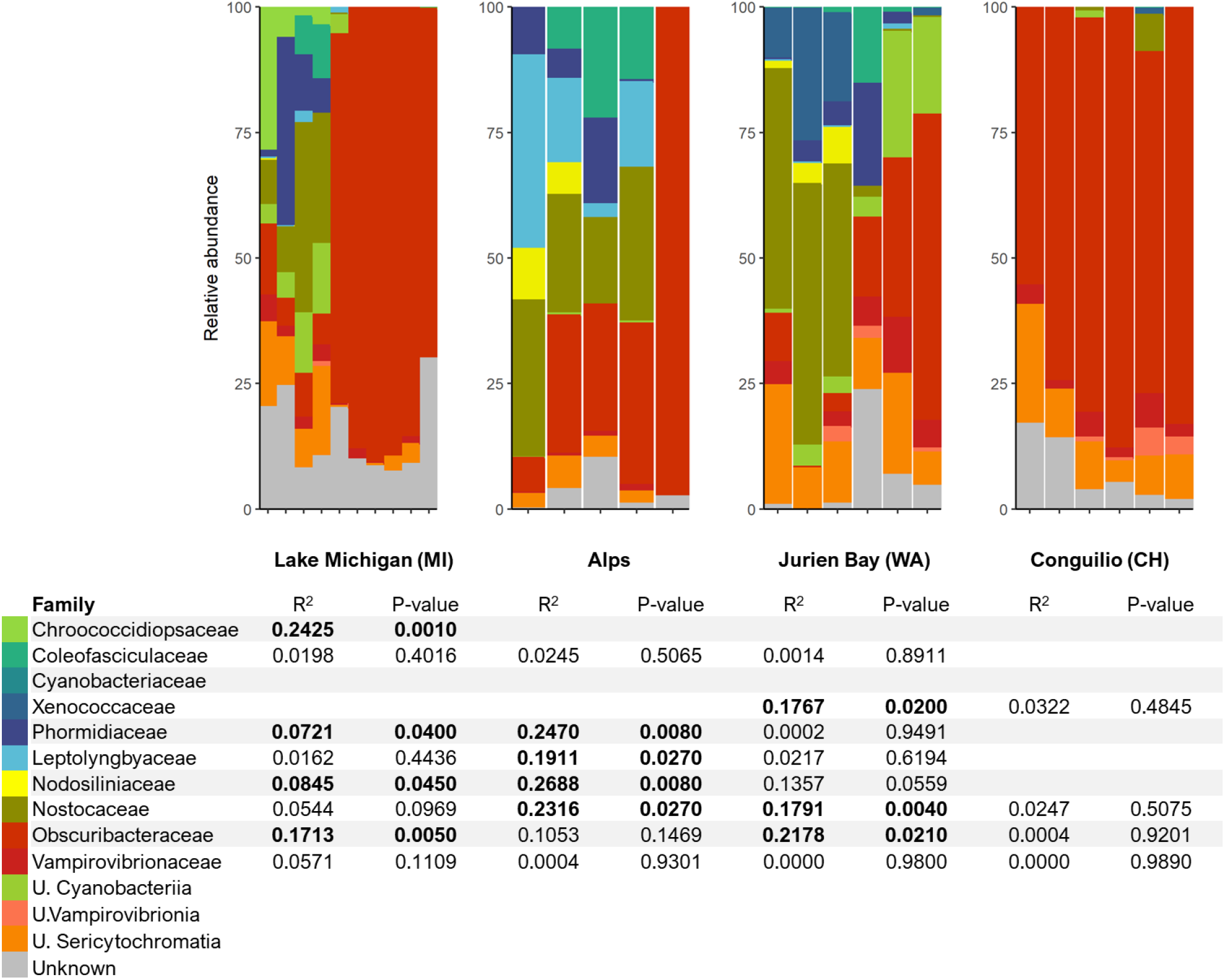
Relative abundance of cyanobacterial families (%) across stages for all chronosequences. Bottom table: Summary of PERMANOVA results for relative abundances of cyanobacterial families in chronosequences with large changes in cyanobacterial composition. U=Unknown.

Changes in the composition of cyanobacterial communities with time occurred in parallel to compositional and structural changes in vegetation. The Alps, Conguilio, Lake Michigan and Jurien Bay chronosequences experienced important changes in vegetation structure and key soil elements (from grasslands to forests; Fig. 3, Fig. S2, Supplementary Table 7). Vegetation shifted from herbaceous to forests in Alps and Lake Michigan, and from cryptogamic covers to shrublands and forests in Conguilio (Fig. 3). Tree and total plant cover had a significant effect on the composition of cyanobacterial communities as soil aged (Supplementary Table 7). The concentration of key soil elements also varied with age in the Alps, Conguilio, Lake Michigan and Jurien Bay chronosequences (Fig. S2), with soil C increasing with soil age in all chronosequences except in Conguilio, as did N and P in the Alps. Soil Nitrogen and available P decreased with soil age in Jurien Bay and Michigan.

### Climate, soil and vegetation determine the structure of soil cyanobacterial communities across ecosystems

Our ordination analyses, which explained a substantial portion of the variance in the cyanobacterial composition (53.5%, see Fig. 5), showed a major impact of vegetation (plant richness, cover of grasses and trees; r^2^=0.37, p<0.01), pH (r^2^=0.32, p<0.01) and MAP (r^2^=0.22, p<0.01) on the composition of soil cyanobacterial communities. The effects of soil age and MAT were not significant. Soil samples were distributed in the PCoA by the presence of forest, confirming that vegetation structure largely regulated the differences in soil cyanobacterial composition across biomes (Fig. 3 and 5).

**Fig.5.**
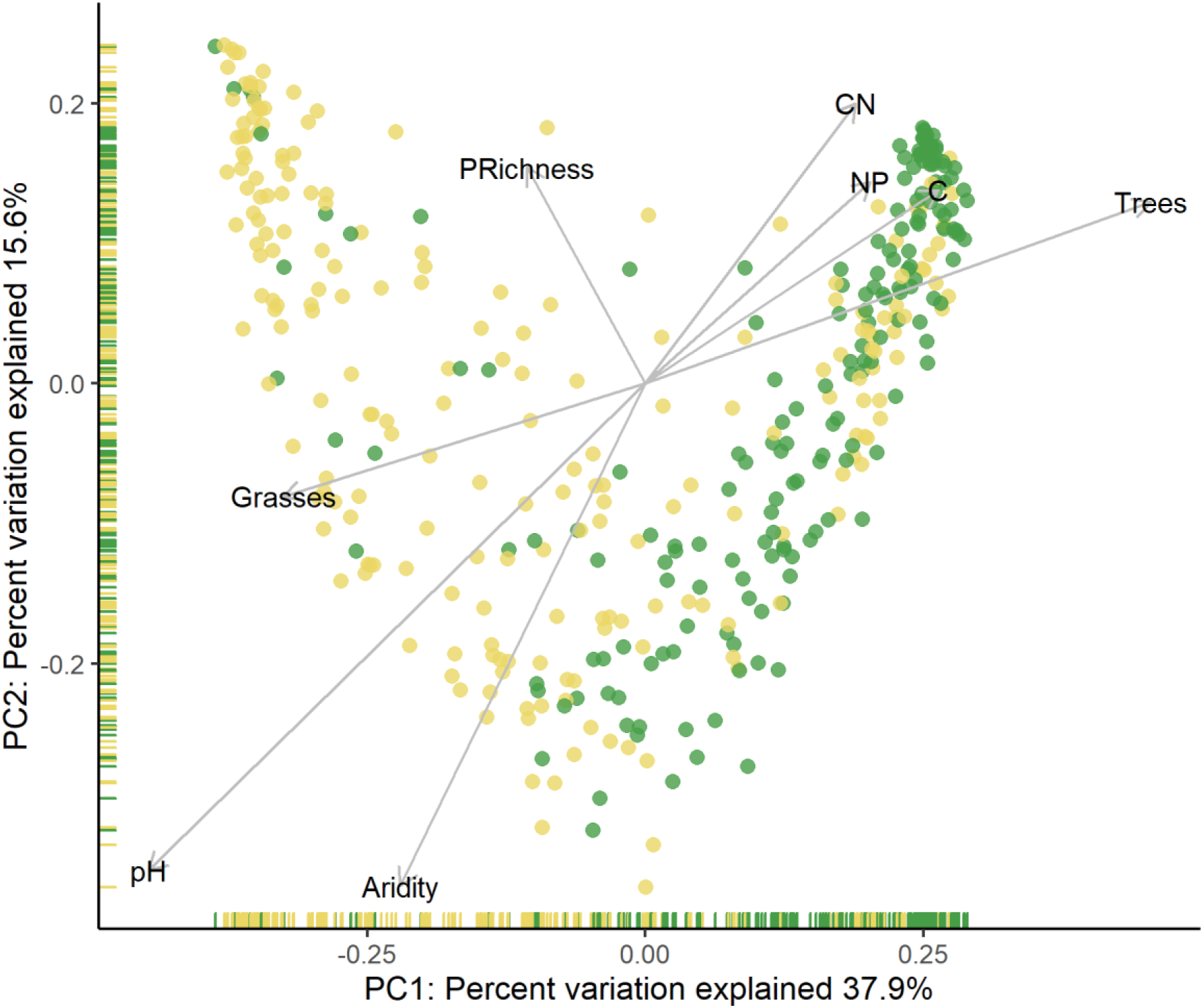
Principal Coordinate Analysis (PCoA) of Unifrac-weighted distances of all chronosequence samples representing forested (green) vs. non forested (yellow) sites. Vectors show important environmental variables, correlated with the PC1 and PC2 axes with corrected Bonferroni p-values <0.05. MAT=Mean Annual Temperature, Soil C= total organic carbon, Soil CN= C:N ratio, Soil NP= N:P ratio, Trees=% tree cover, Grasses= % grass cover.

Structural Equation Models explained 46.6%, 22.4% and 22.8% of the variance in the relative abundances of Vampirovibrionia, Sericytochromatia, Cyanobacteriia, respectively, and 10.6% of the variance in the total abundance of cyanobacteria (Fig. 6 and Fig.S4). Vegetation, soil and climate had strong but differing effects on the total and relative abundance of cyanobacteria. Plant richness and tree cover had negative direct effects on the total abundance of cyanobacteria, and on the relative abundance of Cyanobacteriia and Sericytochromatia. In contrast, tree cover promoted the relative abundance of Vampirovibrionia. However, the presence of herbaceous cover (% grasses cover) had a positive direct effect on relative abundance of Cyanobacteriia and Sericytochromatia. Soil properties and climate also played a key role in regulating cyanobacterial communities. Soil pH had strong and opposing direct effects on the relative abundances of Vampirovibrionia (negative) and Cyanobacteriia (positive). Similar contrasting relationships were found for organic carbon and the relative abundance of Vampirovibrionia (positive) and Cyanobacteriia (negative). The N:P ratio had a direct positive effect on the relative abundance of Cyanobacteriia, and negative effects on those of Sericytochromatia and Vampirovibrionia. Contrasting associations related to climate were also found, as MAP had a positive direct effect on the relative abundance of Cyanobacteriia and negative total effect on the relative abundance of Vampirovibrionia. MAT had a positive direct effect on the relative abundance of Sericytochromatia and a negative total effect on that of Vampirovibrionia. Soil age had a direct negative effect on the relative abundance of Vampirovibrionia and an indirect negative effect on that of Cyanobacteriia via soil pH and total organic C.

**Fig.6:**
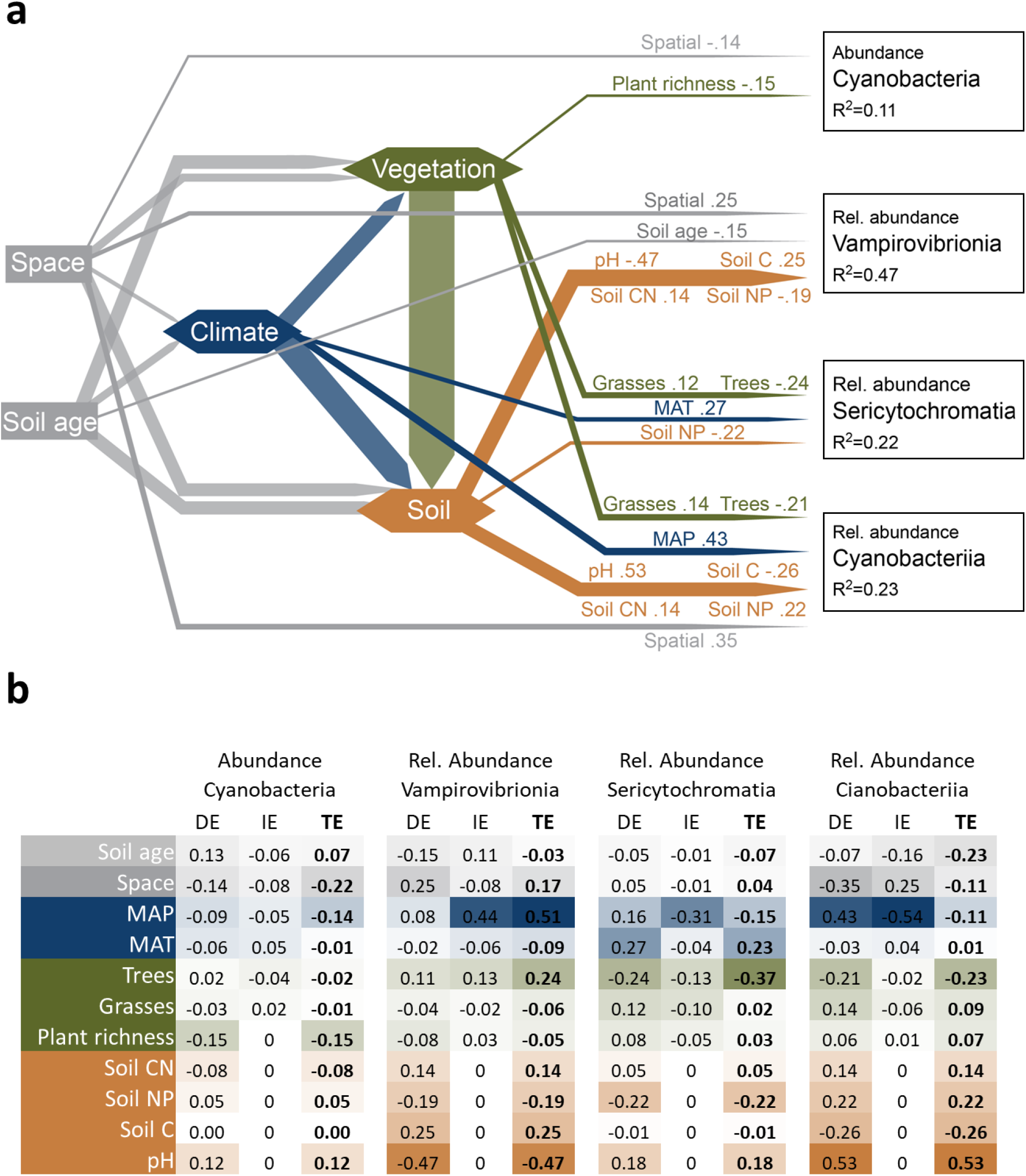
Summary of Structural equation modelling (SEM) results showing the model with direct effects (a) and a table with direct, indirect and total effects (DE, IE, TE) in (b). Variables included are soil age (years), space (Euclidean distances between plots), climate (MAT, and MAP), vegetation (cover of trees and grasses and plant species richness) and soil properties (Soil CN, Soil NP, Soil C and pH) and dependent variables are the abundance of cyanobacteria (measured via qPCR) and the relative abundances of Vampirovibrionia, Sericytochromatia and Cyanobacteriia (Obtained from Miseq, see methods). Numbers in arrows indicate standardized path coefficients. Only significant paths are shown. Hexagonal boxes in the model refer to multiple variables. Model Χ^2^ = 0.375, P= 0.829 df= 2.RMSEA= 0; P= 0.925 The complete SEM model is shown in Fig. S5. MAT=Mean Annual Temperature, Soil C= total organic carbon, Soil CN= C:N ratio, Soil NP= N:P ratio, Trees=% tree cover, Grasses= % grass cover.

## Discussion

Our study provides the first attempt to investigate the changes in soil cyanobacterial communities as soil develops from centuries to millennia across contrasting ecosystems worldwide. We found little effect of soil development *per se* on the development of cyanobacterial communities. Our study suggests that once soil cyanobacteria are established during primary succession under the dominant vegetation, these communities undergo little changes across long periods of time. However, we found cyanobacterial soil communities changes following big vegetational transitions as those from grasslands to forests, so big structural changes in vegetation are key drivers of temporal cyanobacterial community changes.

### Soil cyanobacterial communities do not change during soil development in most of the chronosequences studied

Our first hypothesis, i.e. that soil age would regulate soil cyanobacterial communities, was not fully supported by our data. On the contrary, most chronosequences rather followed our second hypothesis, which suggests that biome type, and in particular vegetation structure, drives the distribution of cyanobacteria worldwide. In most cases, once a cyanobacterial community became established it is generally dominated by members of a particular class (e.g., Cyanobacteriia or Vampirovibrionia) and the community remains stable from centuries to millennia. Previous studies have shown the large influence of climate on soil photosynthetic Cyanobacteriia (Bahl *et al*., 2011; Garcia-Pichel *et al*., 2013; Muñoz-Martín *et al*., 2019). Vegetation and climate are key determinants of the local diversity of Cyanobacteriia, and therefore have a strong influence on their community assembly. On the other hand, dynamics of non-photosynthetic cyanobacteria through space and time are much less known. A study from sediments up to 100 years old in perialpine lakes showed no changes in the community structure of non-photosynthetic cyanobacteria through time, as opposed to the highly changing dynamics of photosynthetic Cyanobacteriia (Monchamp *et al*., 2019). These results suggest that these heterotrophic communities could be shaped by very different environmental drivers than the rest of cyanobacteria through soil development. Here, we observed that Vampirovibrionia dominated across stages in humid temperate forests, and also in the tropical forest of Hawaii and in polar Mount Heckla (Iceland). The presence of canopy (%Tree cover) is beneficial to increment the proportion of this non-photosynthetic group in our soils (positive effects of %Tree cover see Fig 6b) as they don’t depend on solar irradiation and rather depend on organic matter content. These results agree with previous analyses from an independent global survey showing the preference of Vampirovibrionia for humid forests (Cano-Díaz *et al*., 2020). The presence of Vampirovibrionia in all soil chronosequences, and specially their dominance even in first stages in Mount Heckla (Iceland), Chichinautzin (Mexico), Hawaii or the Bolivian chronosequences (Cojiri and Chiar Kkollu), indicates that these cyanobacteria are also colonizers of young soils. Similar patterns have been reported previously for glacial chronosequences (Bardgett *et al*., 2007), where heterotrophic bacterial communities used ancient and recalcitrant carbon as an energy source. On the other side of the ecological spectrum, Cyanobacteriia dominated in arid and semiarid temperate grasslands as they have a preference for areas with low coverage of trees, as seen globally (Garcia-Pichel *et al*., 2003; Bahl *et al*., 2011; Cano-Díaz *et al*., 2020). This way, as tree cover grows the photosynthetic capability of Cyanobacteriia is no longer a beneficial trait and therefore this group loses its dominance (negative effects of %Trees Fig 6b).

During early primary succession (from years to decades) major successional changes in microbial and vegetation structure are expected (Nemergut *et al*., 2007; Liu *et al*., 2016). Previous studies with young chronosequences (up to 10-20 years) showed cyanobacteria increased their abundance and diversity with time (Nemergut *et al*., 2007; Schmidt *et al*., 2008a; Dini-Andreote *et al*., 2014; Liu *et al*., 2016). In contrast, studies with longer time periods (up to 2000 years) showed initial increases in cyanobacterial abundance followed by a steadily decline (Hodkinson *et al*., 2003; Brankatschk *et al*., 2011). Similar patterns were observed in the Merced chronosequence. In Alps and SAGA chronosequences, declines of cyanobacterial abundance and richness, respectively, were observed as soil aged. Conversely, observed increases in cyanobacterial abundance with time in an old chronosequence as Mexico Chichinautzin, and nonlinear responses in Australian Jurien Bay and Chilean Conguilio, are to our knowledge successional patterns not reported before for soil cyanobacteria. These increments could be related to the high proportion of nonphotosynthetic cyanobacteria found in these sites (Fig. 3 and Fig.S5) and the fact that these clades have been recently described. Differences in quantification techniques could explain to some degree the differences between our results and those from previous studies using microscopic techniques or the cover of visible species to assess cyanobacterial abundance (Kastovská *et al*., 2005; Dojani *et al*., 2011). Also, we include relative abundances of difficult-to-culture groups as non-photosynthetic cyanobacteria (Soo *et al*., 2015). Genetic-based quantifications from previous studies from decades or years report steadily declines in cyanobacterial abundance through succession (Kastovská *et al*., 2005; Dini-Andreote *et al*., 2014), pointing out to the need to use both longer time scales and a comprehensive survey, such as those used here, to find novel cyanobacterial successional dynamics. Likewise, the changes in the relative proportion of non-photosynthetic cyanobacteria through soil development reported here are fully novel as they have not been studied before. Nevertheless, in most of the chronosequences studied the abundance and richness of cyanobacteria remained stable over time. These findings have important implications for ecosystem functioning as cyanobacteria enhance soil C and N contents and stability in early ecosystems, paving the way for the development of late-successional communities (Hooper *et al*., 2000). This could provide interesting legacy effects for these soil communities, including the resistance/resilience to environmental changes, all due to an initial colonization that is maintained along soil development (Nemergut *et al*., 2007; Van der Putten *et al*., 2013).

### Shifts in vegetation structure can explain observed changes in soil cyanobacterial communities as soils age

In the chronosequences from Alps, Conguilio, Jurien Bay and Lake Michigan, we observed that changes in vegetation structure from grasslands to forests during ecosystem development were mirrored by changes in the community of cyanobacteria (Fig. 3 and Fig. 4). In these chronosequences, photosynthetic Cyanobacteriia dominated the first chronosequence stages, being replaced by heterotrophic Vampirovibrionia communities as soils developed. These results are probably associated with a change in limiting resources for cyanobacterial communities from carbon in young ecosystems to soil surface availability and light in mature forests (Garcia-Pichel, 2009; Cano-Díaz *et al*., 2019). This agrees with previous studies showing that, after an initial colonization and growth of photosynthetic cyanobacteria, they decline at later successional stages (Hodkinson *et al*., 2003; Brankatschk *et al*., 2011; Dini-Andreote *et al*., 2014). Thus, our results indicate that land use changes from forests to grasslands (e.g., deforestation) would result in important changes in cyanobacterial communities.

The case of Conguilio temperate chronosequence from Chile shows a clear successional effect on vegetation and soil properties, but these were not apparently translated to soil cyanobacterial communities as non-photosynthetic Vampirovibrionia class dominated all chronosequence stages. The ecology of Vampirovibrionia, as the rest of non-photosynthetic cyanobacteria, is still not well known as it has been studied mainly through metagenome-assembled techniques from environmental sequencing, (Di Rienzi *et al*., 2013; Soo *et al*., 2014, 2015, 2017). Vampirovibrionia genomes point to a metabolic capacity for aerobic respiration and a possible adaptation to low oxygen conditions as metagenomes only show C-family oxygen reductases related with microaerobic conditions (Soo *et al*., 2017). Previous research showed community assemblage of non-photosynthetic cyanobacteria remained stable in contrast with Cyanobacteriia which developed seasonal responses to temperature and organic matter content (Monchamp *et al*., 2019). Despite community changes were detected in Conguilio at the Zotu level (PERMANOVAs in Supplementary Table 6), further community analyses showed no evident changes, except for the Sericytochromatia class. The decrease in the relative abundance of Sericytochromatia through soil development is a general pattern observed throughout all the chronosequences (Supplementary Table 4 and S2). This pattern appears to be mediated by vegetation because of the negative influence of tree cover and NP ratio on relative abundance of Sericytochromatia along chronosequences (Fig. 6 and Fig. S4). Little is known about the ecology of Sericytochromatia, as less than ten metagenome assembled sequences are available to date (Soo *et al*., 2017), and further studies are needed before strong generalizations about the mechanisms underlying the results observed can be made.

Changes in the diversity of cyanobacterial communities during the first stages of ecological succession have been previously linked with their functional capacities (e.g., N fixation) (Pointing & Belnap, 2012). Filamentous cyanobacteria such as *Microcoleus vaginatus* are usually the first colonizers of bare soils due to their exopolysaccharide sheath, which make filamentous networks that stabilize soils (Danin *et al*., 1998; Zhang, 2005; Lan *et al*., 2012). Afterwards, heterocystous cyanobacteria such as *Nostoc* sp. and *Scytonema* sp. appear; these species have a highly efficient N fixing capacity through specialized cells (heterocytes) and sometimes contain pigments (e.g. Scytonemin) that protect them from UV radiation, (Danin *et al*., 1998; Hu *et al*., 2004; Lan *et al*., 2015). In Jurien Bay and Alps, we observed a decrease in *Nostocaceae* (heterocystous) abundance with soil age (Fig. 4). In these specific chronosequences, our results rather agree with the classical model of long-term ecosystem development where primary production is limited to N on young soils and P on older ones (Laliberté *et al*., 2012). Cyanobacteria appearing in these first stages have the capacity to fix N (with or without heterocytes), which makes these organisms highly competitive and therefore dominant in the first stages where N is low (Fig. 6). The enrichment in N provided by these cyanobacteria is likely a factor promoting the following colonization by vegetation (Wardle *et al*., 2004; Van der Putten *et al*., 2013). Interestingly, and in contrast with previous reports on changes on the abundance of cyanobacteria at early stages, community turnover were found in old chronosequences (Jurien Bay is 2000 ky, Conguilio is 5000 ky and Alps is 120 ky; only Lake Michigan is relatively young, 4 ky), which shows the benefits of using multiple sites and ages to test and generalize hypotheses about soil microorganisms.

### Concluding remarks

Unlike what has been reported in studies from early ecosystem succession, we did not find strong and significant effects of soil development on the abundance and diversity of cyanobacteria across the global set of chronosequences studied. These cyanobacteria are directly linked with their surrounding vegetation, as we only found profound community changes in those chronosequences wherein vegetation structure has also undergone important changes over time from grasslands to forests. This cyanobacterial compositional shift was mainly related to a turnover from photosynthetic to non-photosynthetic cyanobacteria. These findings significantly advance our knowledge on the ecology and natural history of soil cyanobacteria. This information is relevant to increase our ability to predict future shifts in soil cyanobacteria driven by climate and land use changes.

## Acknowledgements

This project received funding from the European Union’s Horizon 2020 research and innovation program under the Marie Skłodowska-Curie grant agreement 702057 (CLIMIFUN), a Large Research Grant from the British Ecological Society (agreement n° LRA17\1193; MUSGONET), and from the European Research Council (ERC grant agreement n° 647038, BIODESERT). M.D-B. is supported by a Ramón y Cajal grant from the Spanish Government (agreement n° RYC2018-025483-I). C.C-D. acknowledges support from BIODESERT. FTM acknowledges support from Generalitat Valenciana (CIDEGENT/2018/041). We would like to thank the researchers involved in the CLIMIFUN project for the help with soil sampling. We also would like to thank Alberto Benavent-González for his help optimizing the cyanobacterial qPCRs in HIE laboratory of Western Sydney University. The authors declare no conflict of interest.

## Author Contribution

M.D.-B designed research, C.C.-D., F.T.M., J.W., J.L, B.S., V.O., B.G. M.D-B., performed research, C.C.-D. J.W. and M.D-B. analyzed data, C.C.-D, M.D.-B and F.T.M. wrote the paper with all authors contributing to the drafts

