## Supplementary Materials for "Vegetation structure determines cyanobacterial communities during soil development across global biomes"

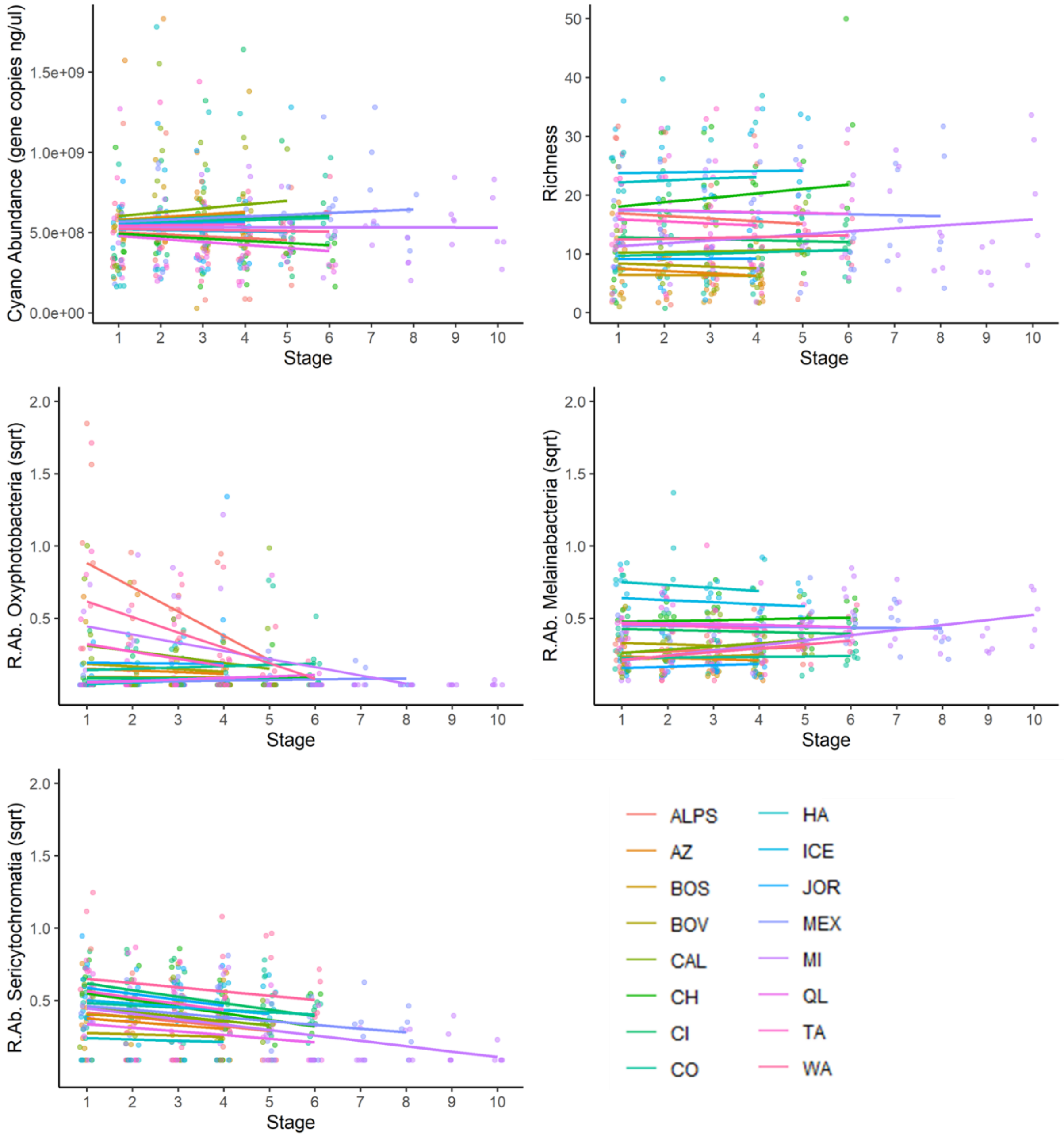


**Figure S1:** Prediction plots of mixed models showing variations of the relationships between chronosequence stage and cyanobacterial abundance, richness and relative abundances of cyanobacterial classes according to random effects (Chronosequence) R. Ab= Relative abundance. See details of the mixed models fitted in Supplementary Table 3.

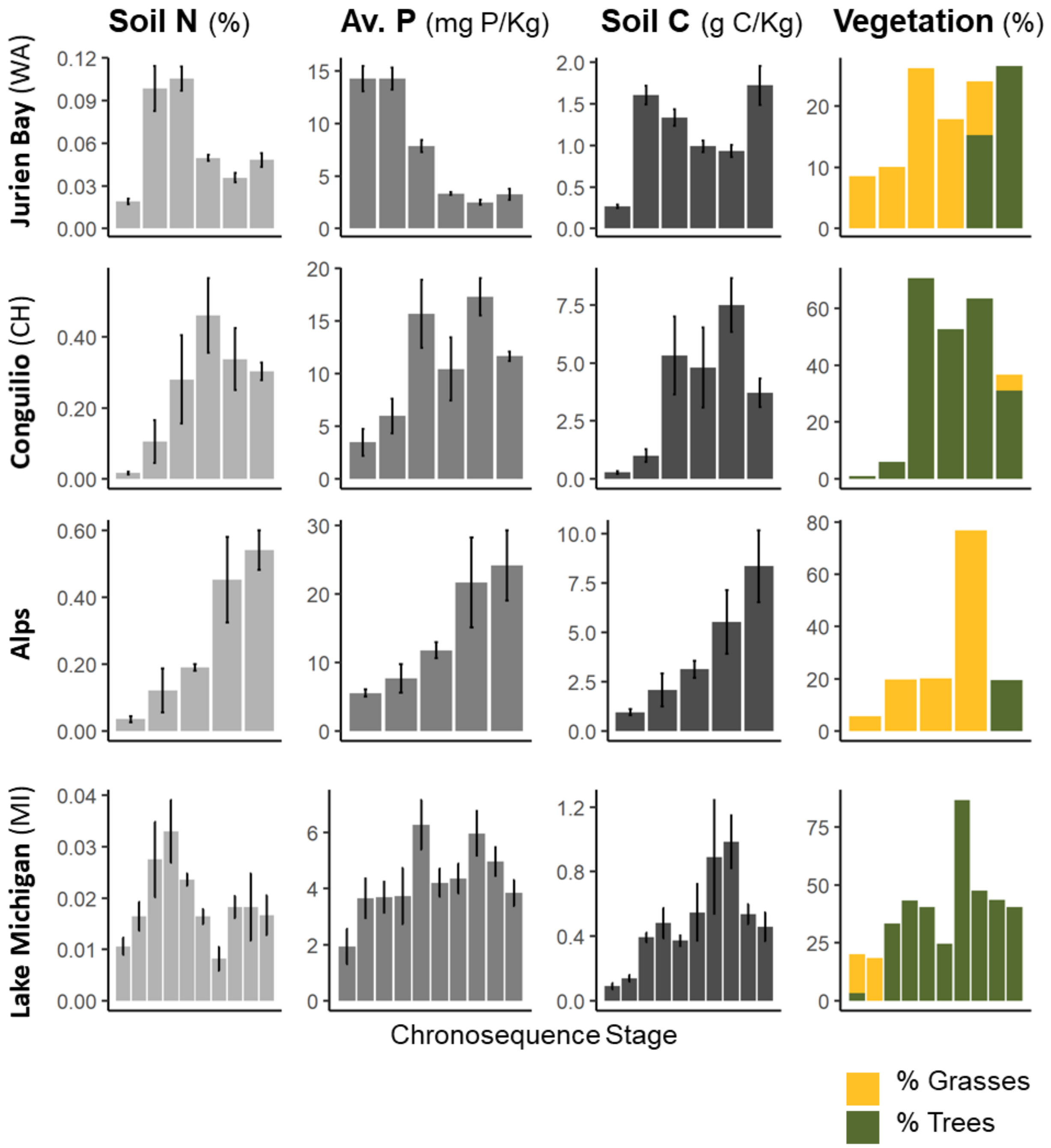


**Figure S2.** Temporal variation across chronosequence stage of key soil (Soil N, Available P [Av.P] and Soil C) and vegetation (cover of Grasses and Trees) variables for selected chronosequences. Data for soil properties are mean ± SD (n =5).

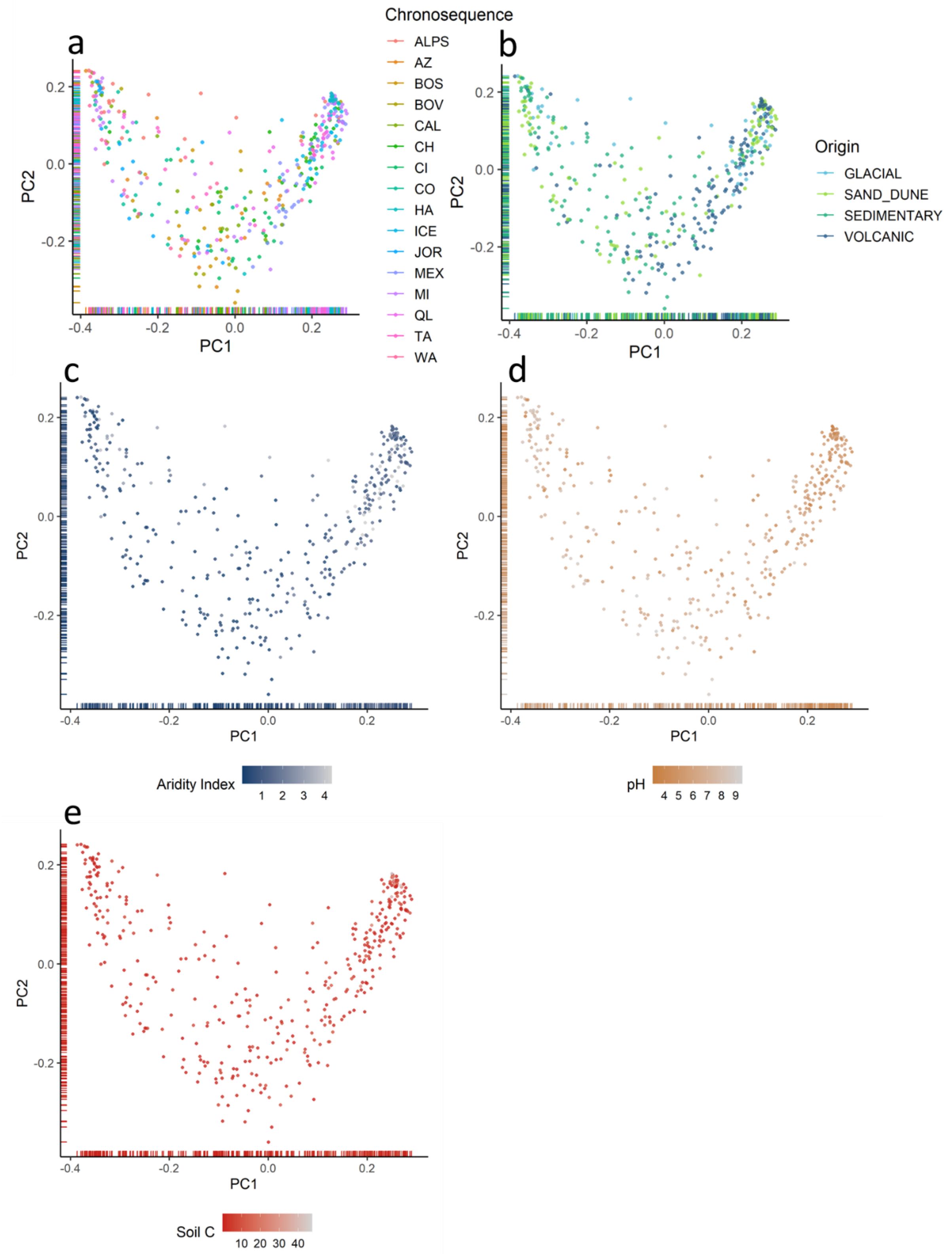


**Figure S3**. Principal Coordinate Analysis (PCoA) of weighted unifrac distances of all chronosequence samples colored by chronosequence (a), chronosequence origin (b), Aridity Index (c), pH (d) and total organic carbon [Soil C] (e).

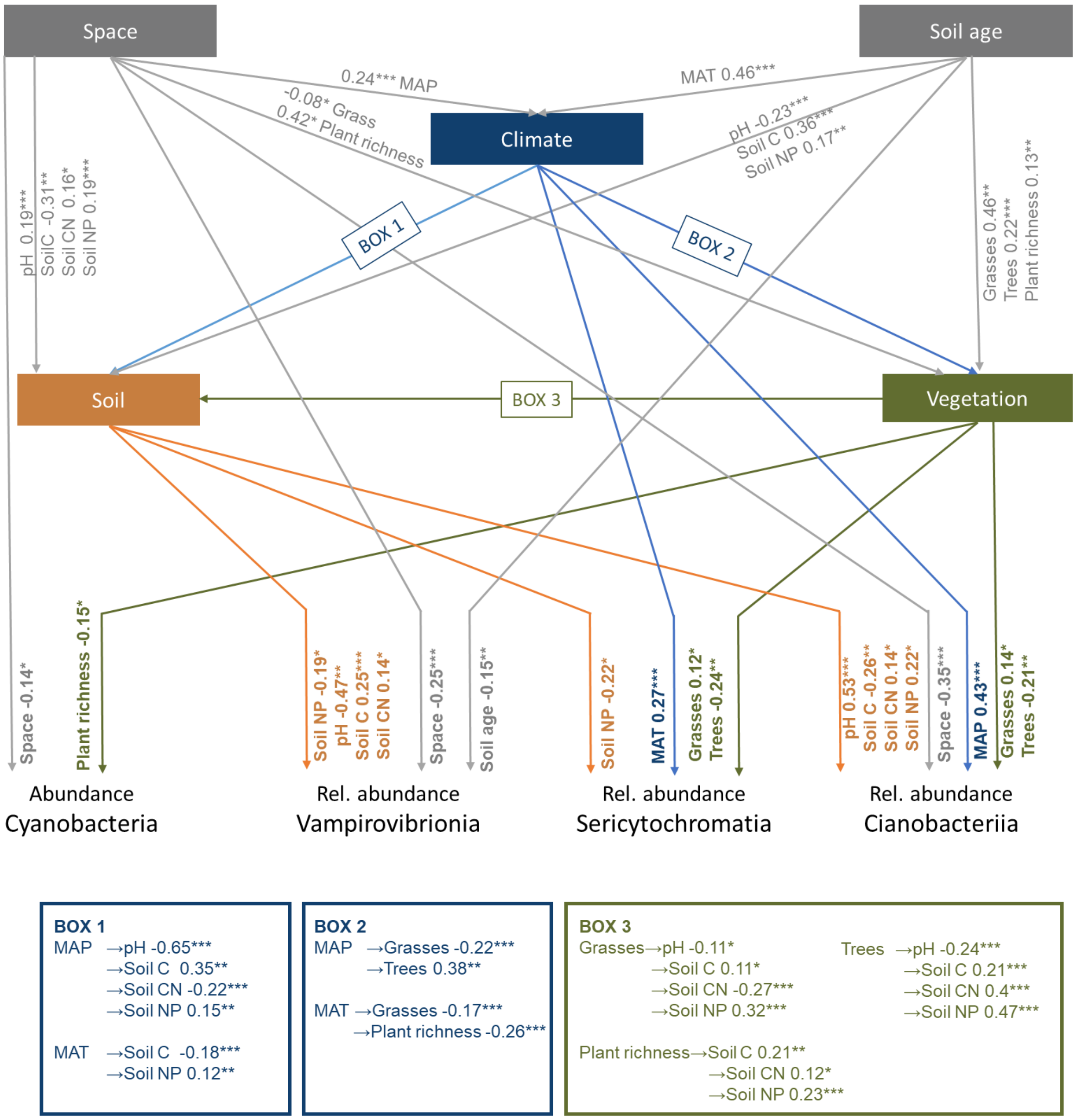


**Figure S4.** Results of the structural equation model used, showing the direct effects of soil age, space (Euclidean distances between plots), climate (MAT, and MAP), vegetation (cover of trees and grasses and plant species richness) and soil properties (Soil CN, Soil NP, Soil C and pH) on the abundance of cyanobacteria and on the relative abundances of Vampirovibrionia, Sericytochromatia and Cyanobacteriia. Numbers in arrows indicate standardized path coefficients. Only significant paths are shown (*P<0.05, **P<0.01, and ***P<0.001). Model $X^2$ =0.375, P= 0.829, df= 2. MAT=Mean Annual Temperature, Soil C= total organic carbon, Soil CN= C:N ratio, Soil NP= N:P ratio, Trees=% tree cover, Grasses= % grass cover.

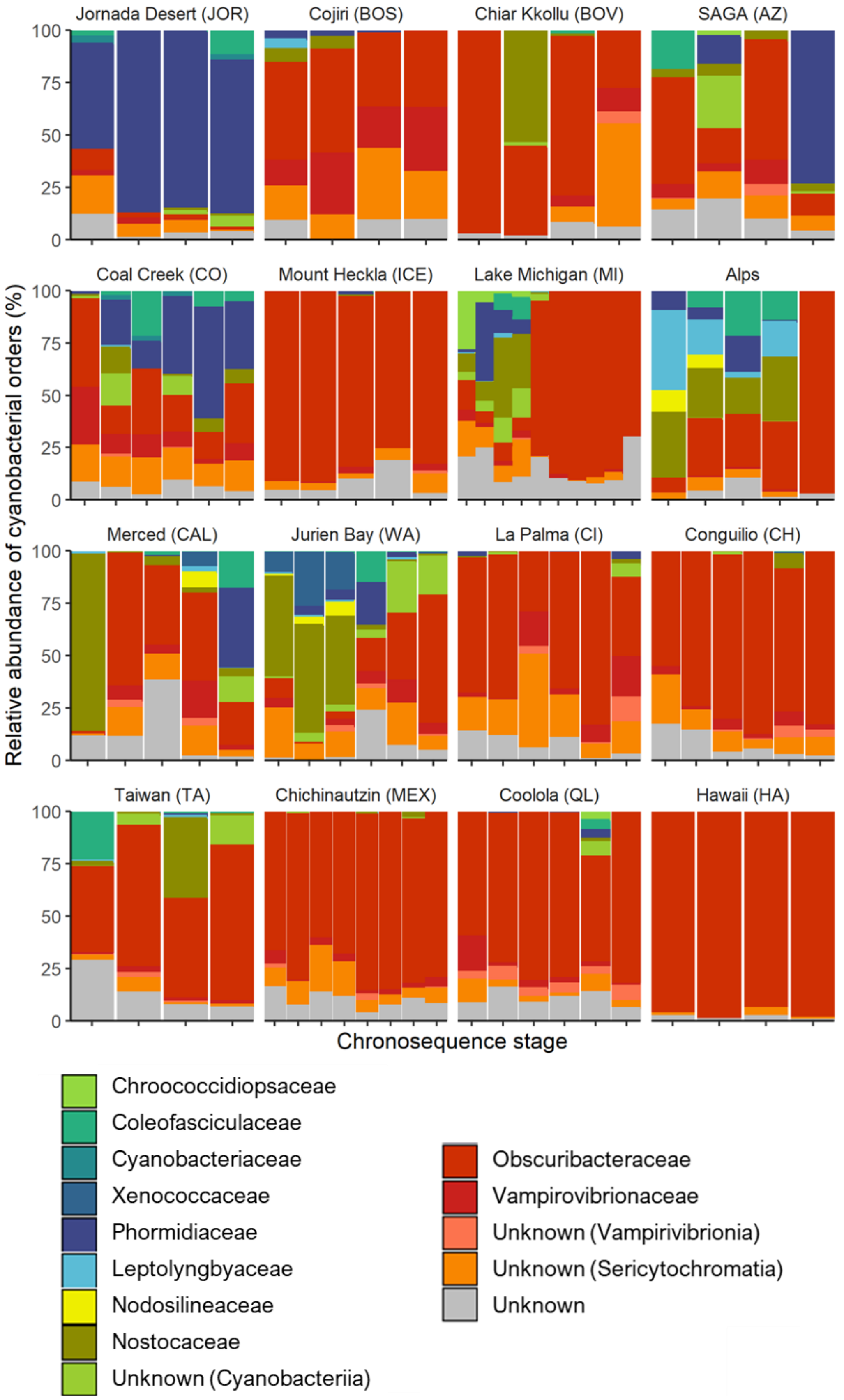


**Figure S5:** Relative abundance (%) of cyanobacterial families across stages for all chronosequences.

**Supplementary Table 1**: List of references from previous works on the chronosequences studied

| | Reference |
|---|---|
| **ALPS** | Kaufmann, R. (2001). Invertebrate Succession on an Alpine Glacier Foreland. Ecology 82, 2261-2278. |
| **AZ** | Selmants, P.C., Hart S.C. (2008). Substrate age and tree islands influence carbon and nitrogen dynamics across a retrogressive semiarid chronosequence. Global Biogeochemical Cycles 22, GB1021. |
| **BOS** | Alfaro, F.D. et al. (2017). Microbial communities in soil chronosequences with distinct parent material: The effect of soil pH and litter quality. Journal of Ecology 105, 1709-1722. |
| **BOV** | Alfaro, F.D. et al. (2017). Microbial communities in soil chronosequences with distinct parent material: The effect of soil pH and litter quality. Journal of Ecology 105, 1709-1722. |
| **CAL** | Doetterl S. et al. (2019). Links among warming, carbon and microbial dynamics mediated by soil mineral weathering. Nature Geoscience 11, 589-593. |
| **CH** | Pérez, C.A. et al. (2017). Biological nitrogen fixation in a post-volcanic chronosequence from south-central Chile. Biogeochemistry 132, 23-36. |
| **CI** | Delgado-Baquerizo M. et al. (2019). Changes in belowground biodiversity during ecosystem development. PNAS 116, 6891-6896. |
| **CO** | Delgado-Baquerizo M. et al. (2019). Changes in belowground biodiversity during ecosystem development. PNAS 116, 6891-6896. |
| **HA** | Chadwick, O.A. et al. (1999). Changing sources of nutrients during four million years of ecosystem development. Nature 397, 491–497. |
| **ICE** | Cutler, N.A. et al. (2008). The spatiotemporal dynamics of a primary succession. Journal of Ecology 96, 231-246. |
| **JOR** | Lajtha K., Schlesinger W.H. (1988). The Biogeochemistry of Phosphorus Cycling and Phosphorus Availability Along a Desert Soil Chronosequence. Ecology 69, 24-39. |
| **MEX** | Peña-Ramírez, V.M. et al. (2015). Rates of pedogenic processes in a chronosequence of volcanic ash soils of Central Mexico. Quaternary International 376, 19-33. |
| **MI** | Williams, M.A. et al. (2013). Bacterial communities in soil mimic patterns of vegetative succession and ecosystem climax but are resilient to change between seasons. Soil Biology and Biochemistry 57, 749-757. |
| **QL** | Walker, J. et al. (2018). Dating the Cooloola coastal dunes of South-Eastern Queensland, Australia. Marine Geology 398, 73–85. |
| **TA** | Tsai, H. et al. (2006). A river terrace soil chronosequence of the pakua tableland in central Taiwan. Soil Science 171, 167-179. |
| **WA** | Laliberté, E. et al. (2014). Environmental filtering explains variation in plant diversity along resource gradients. Science 345, 1602-1605 |

**Supplementary Table 2:** Vegetation and soil age (year estimation) by chronosequence and stage.

| | Stage | Veg. | Dominant vegetation species | Age (yr.) |
|---|---|---|---|---|
| **ALPS** | 1 | Forbland | *Saxifraga azoides, Saxifraga oppositifolia, Poa alpina, Linaria alpina, Artemisia gentipi* | 10 |
| | 2 | Forbland | *Trifolium pallescens, Campanula scheuchzeri, Saxifraga oppositiolia, Saxifraga aizoides* | 45 |
| | 3 | Grassland | *Kobresia myosuroides, Agrostis alpina, Alchemilla fissa, Trifolium pratense spp.* | 125 |
| | 4 | Grassland | *Avenula versicolor, Carex sempervirens, Festuca halleri, Anthoxanthum alpinum* | 10000 |
| | 5 | Forest | *Fagus sylvatica, Abies alba, Acer pseudoplatanus, Picea abies, Quercus robur* | 120000 |
| **AZ** | 1 | Forest | *Juniperus monosperma, Pinus edulis, Bouteloua gracilis* | 900 |
| | 2 | Forest | | 55000 |
| | 3 | Forest | | 750000 |
| | 4 | Forest | | 3000000 |
| **BOS** | 1 | Shrubland | *Astragalus pusillus, Atriplex imbricata, Baccharis boliviensis, Baccharis tola, Ephedra breana, Haplopappus rigidus, Junellia seriphioides, Lycium chanar, Opuntia boliviensis* | 25 |
| | 2 | Shrubland | *Fabiana densa, Atriplex imbricata, Baccharis boliviensis, Lycium chanar, Baccharis tola, Haplopappus rigidus, Hoffmannseggia minor, Junellia seriphioides, Mutisia ledifolia, Nassella curviseta* | 11400 |
| | 3 | Shrubland | *Atriplex imbricata, Baccharis boliviensis, Haplopappus rigidus, Junellia seriphioides, Lycium chanar, Mutisia ledifolia, Nassella curviseta, Trichocereus atacamensis* | 14100 |
| | 4 | Shrubland | *Atriplex imbricata, Baccharis boliviensis, Cheilanthes ternifolia, Diplostephium cinereum, Ephedra breana, Fabiana densa, Lycium chanar, Mutisia ledifolia, Senecio dryophyllus, Senecio nutans, Stevia sp., Trichocereus atacamensis* | 20000 |
| **BOV** | 1 | Shrubland | *Adesmia spinosa, Atriplex imbricata, Chuquiraga atacamensis. Frankenia triandra, Sisymbrium sp., Nassella curviseta* | 25 |
| | 2 | Shrubland | *Opuntia boliviensis, Acantholippia punensis, Atriplex imbricata, Chuquiraga atacamensis, Ephedra breana, Senecio dryophyllus, Sisymbrium sp.* | 11400 |
| | 3 | Shrubland | *Acantholippia punensis, Adesmia spinosa, Atriplex imbricata, Chuquiraga atacamensis, Nassella curviseta* | 14100 |
| | 4 | Shrubland | *Acantholippia punensis, Atriplex imbricata, Chuquiraga atacamensis, Senecio dryophyllus* | 20000 |
| **CAL** | 1 | Shrubland | *Populus fremontii, Helianthus annuus, Amaranthus albus* | 100 |
| | 2 | Shrubland | *Quercus lobata, Silybum marianum, Hordeum murinum L* | 3000 |
| | 3 | Grassland | *Festuca californica* | 30000 |
| | 4 | Grassland | *Rytidosperma penicillatum* | 600000 |
| | 5 | Grassland | *Festuca bromoides, F. myuros, Bromus hordaceous, Bromus diandrus* | 3000000 |
| **CH** | 1 | Moss | *Gaultheria pumila, Racomitrium lanuginosum* | 60 |
| | 2 | Shrubland | *Lomatia hirsuta, Austrocedrus chilensis* | 266 |
| | 3 | Forest | *Araucaria araucana, Nothofagus antarctica* | 776 |
| | 4 | Forest | *Nothofagus dombeyi, Araucaria araucana* | 3470 |
| | 5 | Forest | *Nothofagus dombeyi, N. obliqua, N. alpina* | 60000 |
| | 6 | Forest | *Nothofagus dombeyi, N. alpina* | 5000000 |
| **CI** | 1 | Forest | *Pinus canariensis* | 525 |
| | 2 | Forest | *Pinus canariensis, Erica arborea, Pterocephalus porphyranthus* | 6000 |
| | 3 | Forest | *Pinus canariensis, Adenocarpus viscosus, Chamaecytisus proliferus, Erica arborea* | 40000 |
| | 4 | Forest | *Pinus canariensis, Adenocarpus viscosus* | 600000 |
| | 5 | Forest | *Pinus canariensis, Cistus symphytifolius* | 1100000 |
| | 6 | Forest | *Pinus canariensis, Cistus symphytifolius* | 1700000 |
| **CO** | 1 | Grassland | *Juncus arcticus, Andropogon gerardii, Panicum virgatum* | 5000 |
| | 2 | Grassland | *Andropogon gerardii, Panicum virgatum* | 140000 |
| | 3 | Grassland | *Panicum virgatum, Poa compresa, Andropogon gerardii* | 240000 |
| | 4 | Grassland | *Chrysopsis sp., Andropogon gerardii, L. cinquefoil* | 640000 |
| | 5 | Grassland | *Andropogon gerardii, M. Burgia, Poa compresa,* | 1000000 |
| | 6 | Grassland | *Andropogon gerardii, Poa compresa, M. Burgia* | 2000000 |
| **HA** | 1 | Forest | *Metrosideros polymorpha, Morella faya, Vaccinium calycinum, Ilex anomala, Cheirodendron trigynum, Cibotium glaucom, Hedychium gardnerianum, Isoetes sp. (grass), Coprosma sp., Myrsine lessertiana, Dicranopteris linearis, Machaerina angustifolia, Anemone hupehensis,* | 300 |
| | 2 | Forest | *Metrosideros polymorpha, Cheirodendron trigynum, Cibotium glaucom, Cibotium menziesii, Ilex anomala, Freycinetia arborea, Astelia menziesii, Melicope clusiifolia, Vaccinium calycinum, Nephrolepis sp., Asplenium spp. (multi), Athyrium microphyllum, Ilex myrtifolia, Peperomia sp., Polypodium sp.,* | 20000 |
| | 3 | Forest | *Metrosideros polymorpha, Cibotium glaucom, Cibotium menziesii, Hedychium gardnerianum, Vaccinium calycinum, Cheirodendron trigynum, Psidium cattleianum, Dicranopteris linearis, Asplenium sp., Melicope clusiifolia, Myrsine sandwicensis,* | 150000 |

| | | | | |
|---|---|---|---|---|
| | | | *Elaphoglossum sp., Polygonum punctatum Tibouchina herbacea, Peperomia sp., Psilotum nudum* | |
| | 4 | Forest | *Metrosideros polymorpha, Hedychium gardnerianum, Dicranopteris linearis, Pittosporum gayanum, Psidium cattleianum, Astelia menziesiana, Morella faya, Vaccinium meyenianum, Smilax hawaiensis, Elaphoglossum spp, Elaeocarpus bifidus, Clerodendrum sp., Alyxia oliviformis,* | 4100000 |
| **ICE** | 1 | Moss | *Racomitrium lanuginosum; Empetrum nigrum; Stereocaulon vesuvianum* | 172 |
| | 2 | Moss | *Racomitrium lanuginosum; Empetrum nigrum; Arctostaphylos uva-ursi* | 463 |
| | 3 | Moss | *Racomitrium lanuginosum; Betula nana; Hylocomium splendens; Empetrum nigrum; Arctostaphylos uva-ursi* | 628 |
| | 4 | Moss | *Racomitrium lanuginosum; Salix phylicifolia; Empetrum nigrum; Hylocomium splendens* | 717 |
| | 5 | Shrubland | *Racomitrium lanuginosum; Empetrum nigrum; Betula nana; Hylocomium splendens* | 859 |
| **JOR** | 1 | Forbland | *Opuntia phaeacantha. var., Boerhavia spp., Eragrostis lehmanniana.* | 1100 |
| | 2 | Grassland | *Sporobolus contractus, Muhlenbergia porteri, Larrea tridentata, Ephedra trifurca.* | 2200 |
| | 3 | Forbland | *Boerhavia spp., Larrea tridentata .* | 8000 |
| | 4 | Forbland | *Boerhavia spp., Ephedra trifurca Torr., Erioneuron pulchellum* | 25000 |
| **MEX** | 1 | Forest | *Pinus montezumae, Bacharis conferta, Alnus firmifolia, Penstemon sp.* | 1000 |
| | 2 | Forest | *Abies religiosa, Arbutus xalapensis, Pinus herrerae, Bacharis conferta, Pinus montezumae, Penstemon sp., Bacharis conferta* | 1835 |
| | 3 | Forest | *Pinus montezumae, Pinus pseudostrobus, Alnus firmifolia, Quercus laurina,* | 3800 |
| | 4 | Forest | *Pinus montezumae, Alnus firmifolia* | 6200 |
| | 5 | Forest | *Pinus montezumae, Bacharis conferta, Buddleja parviflora* | 8000 |
| | 6 | Forest | *Pinus patula, Alnus firmifolia, Pinus montezumae, Senecio sp.* | 10000 |
| | 7 | Forest | *Pinus ayacahuite, Pinus pseudostrobus, Pinus montezumae* | 30500 |
| | 8 | Forest | *Pinus montezumae, Abies religiosa, Quercus laurina, Penstemos sp., Bacharis conferta* | 100000 |
| **MI** | 1 | Grassland | *Amophilous breviligulata, Agropyron dasystachium, Cerisium pitheri, Arctostaphylos uva-ursi* | 73 |
| | 2 | Grassland | *Amophilous breviligulata, Agropyron dasystachium, Cerisium pitheri, Arctostaphylos uva-ursi, Schizachyrium scoparium* | 113 |
| | 3 | Shrubland | *Arctostaphylos uva-ursi, Juniperus communis, Pinus strobus* | 163 |
| | 4 | Shrubland | *Pteridium aquilinum, Pinus resinosa, Abies sp.* | 243 |
| | 5 | Forest | *Gaultheria procumbens, Pinus resinosa,* | 485 |
| | 6 | Forest | *Abies balsamea, Pinus resinosa, Juniperus communis* | 863 |
| | 7 | Forest | *Abies balsamea, Pinus resinosa, Pinus strobus* | 1400 |
| | 8 | Forest | *Pinus resinosa, Vaccinium myrtilloides, Gaultheria procumbens* | 2500 |
| | 9 | Forest | *Pinus resinosa, Vaccinium myrtilloides, Gaultheria procumbens* | 3200 |
| | 10 | Forest | *Pinus resinosa, Pinus strobus, Gaultheria procumbens* | 4000 |
| **QL** | 1 | Forest | *Eucalyptus tessellaris, Angophora costata, Eucalyptus intermedia, Casuarina littoralis, Melaleuca quinquenerva, Banksia integrifolia, Banksia serrata, Macrozamia spp., Acacia aulacocarpa, Acacia flavescens, Cassytha paniculata, Gahnia sieberiana,* | 3600 |
| | 2 | Forest | | 6700 |
| | 3 | Forest | | 134000 |
| | 4 | Forest | | 176000 |
| | 5 | Forest | | 324000 |
| | 6 | Forest | | 716000 |
| **TA** | 1 | Shrubland | *Tea camellia* | 28000 |
| | 2 | Shrubland | | 105000 |
| | 3 | Shrubland | | 322000 |
| | 4 | Shrubland | | 399000 |
| **WA** | 1 | Shrubland | *Acacia cyclops, Acacia rostellifera, Scaevola crassifolia, Olearia axillaris, Spyridium globulosum* | 100 |
| | 2 | Shrubland | *Melaleuca systena, Acacia lasiocarpa, Acacia rostellifera* | 1000 |
| | 3 | Shrubland | *Melaleuca systena, Acacia lasiocarpa, Acacia rostellifera* | 6500 |
| | 4 | Shrubland | *Melaleuca systena, Banksia leptophylla, Calothamnus quadrifidus* | 120000 |
| | 5 | Shrubland | *Banksia menziesii, Banksia attenuata, Mesomelaena pseudostygia, Hibbertia hypericoides* | 480000 |
| | 6 | Shrubland | *Banksia menziesii, Jacksonia floribunda, Banksia leptophylla* | 2000000 |

**Supplementary Table 3**: Mixed models with random intercept and slope fitted for abundance, richness and relative abundance of the main cyanobacterial classes. Significant models (p<0.05) are in bold.

| | Fixed Effects | Sum Sq | Mean Sq | NumDF | DenDF | F | P-value |
|---|---|---|---|---|---|---|---|
| **Abundance Cyanobacteria** | Stage | 8.71E+15 | 8.71E+15 | 1 | 11.1990 | 0.1177 | 0.7379 |
| **Richness Phylotypes** | Stage | 9.00E-02 | 9.00E-02 | 1 | 7.7131 | 0.0018 | 0.9676 |
| **Rel ab. Sericytochromatia** | **Stage** | **5.07E-01** | **5.07E-01** | **1** | **5.6396** | **13.0880** | **0.0124** |
| **Rel ab. Vampirovibrionia** | Stage | 2.46E-03 | 2.46E-03 | 1 | 12.8230 | 0.1194 | 0.7352 |
| **Rel ab. Cianobacteriia** | Stage | 1.30E-01 | 1.30E-01 | 1 | 11.7270 | 2.8206 | 0.1195 |

*t-tests use Satterthwaite's method [lmerModLmerTest]*
*Formula: lmer(Variable ~ Stage + (1 + Stage|Chronosequence)*

**Supplementary Table 4.** Linear and polynomial models of cyanobacterial abundance through soil development for all chronosequences. Significant ($p<0.05$) models in bold

| Chronosequence | Abundance Models | DF | Adj- $R^2$ | F | p-value | AIC |
|---|---|---|---|---|---|---|
| ALPS | Linear Model | 17 | 0.0847 | 2.6646 | 0.1210 | 394.8557 |
| | Quadratic Model | 16 | 0.0477 | 1.4506 | 0.2637 | 396.4562 |
| | Cubic Model | 15 | 0.1115 | 1.7530 | 0.1992 | 395.9119 |
| AZ | Linear Model | 16 | -0.0620 | 0.0077 | 0.9310 | 374.7171 |
| | Quadratic Model | 15 | -0.1002 | 0.2256 | 0.8007 | 376.1923 |
| | Cubic Model | 14 | -0.1670 | 0.1892 | 0.9020 | 378.0106 |
| BOS | Linear Model | 17 | -0.0586 | 0.0031 | 0.9561 | 390.1572 |
| | Quadratic Model | 16 | -0.0866 | 0.2828 | 0.7573 | 391.5005 |
| | Cubic Model | 15 | 0.1221 | 1.8346 | 0.1842 | 388.2220 |
| BOV | Linear Model | 7 | -0.1086 | 0.2162 | 0.6561 | 175.0001 |
| | Quadratic Model | 6 | -0.2426 | 0.2192 | 0.8093 | 176.6393 |
| | Cubic Model | 5 | -0.4614 | 0.1581 | 0.9201 | 178.4583 |
| **CAL** | Linear Model | 17 | -0.0556 | 0.0518 | 0.8227 | 393.3992 |
| | **Quadratic Model** | **16** | **0.3114** | **5.0692** | **0.0197** | **386.1314** |
| | **Cubic Model** | **15** | **0.5307** | **7.7836** | **0.0023** | **379.6211** |
| CH | Linear Model | 26 | 0.0109 | 1.2963 | 0.2653 | 549.7652 |
| | Quadratic Model | 25 | 0.0071 | 1.0963 | 0.3496 | 550.7735 |
| | Cubic Model | 24 | -0.0303 | 0.7353 | 0.5412 | 552.6654 |
| CI | Linear Model | 22 | -0.0429 | 0.0548 | 0.8170 | 491.2421 |
| | Quadratic Model | 21 | 0.0331 | 1.3941 | 0.2701 | 490.3099 |
| | Cubic Model | 20 | -0.0046 | 0.9648 | 0.4287 | 492.0582 |
| **CO** | Linear Model | 26 | -0.0136 | 0.6368 | 0.4321 | 548.8015 |
| | Quadratic Model | 25 | -0.0424 | 0.4510 | 0.6421 | 550.4867 |
| | **Cubic Model** | **24** | **0.1831** | **3.0179** | **0.0495** | **544.5169** |
| HA | Linear Model | 10 | 0.2513 | 4.6926 | 0.0555 | 255.7455 |
| | Quadratic Model | 9 | 0.1715 | 2.1384 | 0.1738 | 257.6971 |
| | Cubic Model | 8 | 0.1636 | 1.7173 | 0.2403 | 258.3972 |
| ICE | Linear Model | 20 | -0.0465 | 0.0665 | 0.7991 | 442.8571 |
| | Quadratic Model | 19 | 0.0195 | 1.2084 | 0.3206 | 442.2960 |
| | Cubic Model | 18 | -0.0300 | 0.7959 | 0.5121 | 444.1897 |
| JOR | Linear Model | 17 | -0.0286 | 0.5001 | 0.4890 | 380.5904 |
| | Quadratic Model | 16 | -0.0862 | 0.2859 | 0.7551 | 382.4741 |
| | Cubic Model | 15 | -0.0371 | 0.7851 | 0.5206 | 382.3701 |
| **MEX** | **Linear Model** | **32** | **0.1481** | **6.7374** | **0.0141** | **673.9656** |
| | **Quadratic Model** | **31** | **0.1568** | **4.0693** | **0.0270** | **674.5358** |
| | Cubic Model | 30 | 0.1383 | 2.7656 | 0.0590 | 676.1603 |
| MI | Linear Model | 40 | -0.0127 | 0.4877 | 0.4890 | 821.6124 |
| | Quadratic Model | 39 | -0.0347 | 0.3130 | 0.7331 | 823.4526 |
| | Cubic Model | 38 | -0.0156 | 0.7895 | 0.5073 | 823.5820 |
| QL | Linear Model | 25 | -0.0151 | 0.6135 | 0.4408 | 536.9364 |
| | Quadratic Model | 24 | -0.0458 | 0.4303 | 0.6553 | 538.6398 |
| | Cubic Model | 23 | -0.0275 | 0.7684 | 0.5235 | 539.0121 |
| TA | Linear Model | 16 | 0.0636 | 2.1542 | 0.1616 | 368.7707 |
| | Quadratic Model | 15 | 0.0673 | 1.6130 | 0.2320 | 369.5379 |
| | Cubic Model | 14 | 0.0165 | 1.0951 | 0.3837 | 371.2500 |
| **WA** | **Linear Model** | **24** | **0.1981** | **7.1748** | **0.0131** | **503.7612** |
| | **Quadratic Model** | **23** | **0.1861** | **3.8585** | **0.0359** | **505.0393** |
| | Cubic Model | 22 | 0.1893 | 2.9453 | 0.0553 | 505.7832 |

**Supplementary Table 5.** Linear and polynomial models of cyanobacterial richness through soil development for all chronosequences. Significant (p<0.05) models in bold.

| Chronosequence | Richness Models | DF | Adj- $R^2$ | F | p-value | AIC |
|---|---|---|---|---|---|---|
| ALPS | Linear Model | 20 | 0.0193 | 1.4133 | 0.2484 | 82.9069 |
| | Quadratic Model | 19 | -0.0086 | 0.9105 | 0.4192 | 84.3955 |
| | Cubic Model | 18 | 0.0702 | 1.5281 | 0.2414 | 83.4176 |
| **AZ** | **Linear Model** | **17** | **0.1964** | **5.3992** | **0.0328** | **48.6659** |
| | **Quadratic Model** | **16** | **0.2493** | **3.9888** | **0.0393** | **48.22023** |
| | Cubic Model | 15 | 0.2018 | 2.5166 | 0.0976 | 50.1606 |
| BOS | Linear Model | 17 | -0.0571 | 0.0278 | 0.8695 | 45.8196 |
| | Quadratic Model | 16 | -0.0994 | 0.1864 | 0.8317 | 47.4131 |
| | Cubic Model | 15 | -0.1059 | 0.4257 | 0.7374 | 48.2983 |
| BOV | Linear Model | 7 | -0.0317 | 0.7542 | 0.4139 | 31.5196 |
| | Quadratic Model | 6 | -0.1981 | 0.3387 | 0.7255 | 33.4779 |
| | Cubic Model | 5 | -0.4351 | 0.1915 | 0.8979 | 35.4619 |
| CAL | Linear Model | 23 | -0.0125 | 0.7026 | 0.4105 | 59.5364 |
| | Quadratic Model | 22 | -0.0370 | 0.5717 | 0.5727 | 61.0219 |
| | Cubic Model | 21 | -0.0419 | 0.6785 | 0.5749 | 61.9758 |
| **CH** | Linear Model | 27 | 0.0724 | 3.1854 | 0.0855 | 92.2698 |
| | Quadratic Model | 26 | 0.0698 | 2.0503 | 0.1490 | 93.2570 |
| | **Cubic Model** | **25** | **0.4108** | **7.5070** | **0.0010** | **80.8773** |
| CI | Linear Model | 27 | -0.0338 | 0.0850 | 0.7729 | 79.5939 |
| | Quadratic Model | 26 | -0.0587 | 0.2232 | 0.8015 | 81.1914 |
| | Cubic Model | 25 | -0.1008 | 0.1450 | 0.9319 | 83.1847 |
| CO | Linear Model | 27 | 0.0496 | 2.4608 | 0.1284 | 68.3146 |
| | Quadratic Model | 26 | 0.0162 | 1.2302 | 0.3087 | 70.2222 |
| | Cubic Model | 25 | -0.0182 | 0.8334 | 0.4882 | 72.0798 |
| HA | Linear Model | 11 | -0.0542 | 0.3828 | 0.5487 | 48.7909 |
| | Quadratic Model | 10 | -0.1383 | 0.2711 | 0.7680 | 50.5492 |
| | Cubic Model | 9 | 0.2191 | 2.1222 | 0.1676 | 46.2810 |
| ICE | Linear Model | 23 | -0.0409 | 0.0573 | 0.8130 | 57.0699 |
| | Quadratic Model | 22 | 0.0347 | 1.4312 | 0.2604 | 56.0742 |
| | Cubic Model | 21 | 0.0178 | 1.1447 | 0.3541 | 57.3457 |
| JOR | Linear Model | 18 | -0.0517 | 0.0664 | 0.7996 | 49.0172 |
| | Quadratic Model | 17 | -0.0539 | 0.5145 | 0.6068 | 49.9156 |
| | Cubic Model | 16 | -0.0467 | 0.7176 | 0.5559 | 50.5662 |
| MEX | Linear Model | 37 | -0.0023 | 0.9144 | 0.3452 | 111.2801 |
| | Quadratic Model | 36 | -0.0299 | 0.4480 | 0.6424 | 113.2734 |
| | Cubic Model | 35 | -0.0284 | 0.6506 | 0.5879 | 114.1158 |
| MI | Linear Model | 46 | 0.0507 | 3.5083 | 0.0674 | 145.1623 |
| | Quadratic Model | 45 | 0.0631 | 2.5822 | 0.0868 | 145.4753 |
| | Cubic Model | 44 | 0.0960 | 2.6638 | 0.0595 | 144.6796 |
| QL | Linear Model | 27 | -0.0099 | 0.7265 | 0.4015 | 78.5329 |
| | Quadratic Model | 26 | 0.0340 | 1.4934 | 0.2432 | 78.1493 |
| | Cubic Model | 25 | -0.0042 | 0.9609 | 0.4266 | 80.1381 |
| TA | Linear Model | 17 | 0.0829 | 2.6272 | 0.1234 | 62.2313 |
| | Quadratic Model | 16 | 0.0995 | 1.9944 | 0.1685 | 62.7326 |
| | Cubic Model | 15 | 0.2144 | 2.6373 | 0.0876 | 60.9131 |
| **WA** | Linear Model | 24 | -0.0127 | 0.6861 | 0.4157 | 68.6141 |
| | Quadratic Model | 23 | 0.1156 | 2.6341 | 0.0933 | 65.9846 |
| | **Cubic Model** | **22** | **0.3938** | **6.4124** | **0.0028** | **57.01123** |

**Supplementary Table 6.** PERMANOVA analysis with Unifrac distances for each chronosequence. Chronosequences with large community changes ($p<0.01$) are in bold

| Cronosequence | | Df | SS | MS | Pseudo F | $R^2$ | P-value |
|---|---|---|---|---|---|---|---|
| **ALPS** | **Stage** | **1** | **0.4072** | **0.4072** | **4.5781** | **0.1863** | **0.0110** |
| | **Residuals** | **20** | **1.7787** | **0.0889** | **0.8137** | | |
| | **Total** | **21** | **2.1859** | **1.0000** | | | |
| AZ | Stage | 1 | 0.1768 | 0.1768 | 1.3646 | 0.0743 | 0.2206 |
| | Residuals | 17 | 2.2028 | 0.1296 | 0.9257 | | |
| | Total | 18 | 2.3796 | 1.0000 | | | |
| BOS | Stage | 1 | 0.0968 | 0.0968 | 1.0486 | 0.0581 | 0.3781 |
| | Residuals | 17 | 1.5687 | 0.0923 | 0.9419 | | |
| | Total | 18 | 1.6655 | 1.0000 | | | |
| BOV | Stage | 1 | 0.3314 | 0.3314 | 1.5332 | 0.1797 | 0.1319 |
| | Residuals | 7 | 1.5131 | 0.2162 | 0.8203 | | |
| | Total | 8 | 1.8445 | 1.0000 | | | |
| CAL | Stage | 1 | 0.2344 | 0.2344 | 1.9614 | 0.0786 | 0.0741 |
| | Residuals | 23 | 2.7491 | 0.1195 | 0.9214 | | |
| | Total | 24 | 2.9835 | 1.0000 | | | |
| **CH** | **Stage** | **1** | **0.1735** | **0.1734** | **2.8072** | **0.0942** | **0.0131** |
| | **Residuals** | **27** | **1.6682** | **0.0618** | **0.9058** | | |
| | **Total** | **28** | **1.8417** | **1.0000** | | | |
| CI | Stage | 1 | 0.1781 | 0.1781 | 1.7517 | 0.0609 | 0.0733 |
| | Residuals | 27 | 2.7457 | 0.1017 | 0.9391 | | |
| | Total | 28 | 2.9238 | 1.0000 | | | |
| CO | Stage | 1 | 0.1791 | 0.1791 | 2.7156 | 0.0914 | 0.0218 |
| | Residuals | 27 | 1.7806 | 0.0659 | 0.9086 | | |
| | Total | 28 | 1.9597 | 1.0000 | | | |
| HA | Stage | 1 | 0.0326 | 0.0326 | 0.7433 | 0.0633 | 0.5048 |
| | Residuals | 11 | 0.4831 | 0.0439 | 0.9367 | | |
| | Total | 12 | 0.5157 | 1.0000 | | | |
| ICE | Stage | 1 | 0.0478 | 0.0478 | 1.3893 | 0.0570 | 0.1652 |
| | Residuals | 23 | 0.7918 | 0.0344 | 0.9430 | | |
| | Total | 24 | 0.8396 | 1.0000 | | | |
| JOR | Stage | 1 | 0.2328 | 0.2327 | 2.9612 | 0.1413 | 0.0546 |
| | Residuals | 18 | 1.4148 | 0.0786 | 0.8587 | | |
| | Total | 19 | 1.6475 | 1.0000 | | | |
| MEX | Stage | 1 | 0.0951 | 0.0951 | 1.4693 | 0.0382 | 0.1390 |
| | Residuals | 37 | 2.3959 | 0.0648 | 0.9618 | | |
| | Total | 38 | 2.4911 | 1.0000 | | | |
| **MI** | **Stage** | **1** | **2.3315** | **2.3315** | **26.7790** | **0.3680** | **0.0001** |
| | **Residuals** | **46** | **4.0050** | **0.0871** | **0.6321** | | |
| | **Total** | **47** | **6.3365** | **1.0000** | | | |
| QL | Stage | 1 | 0.1301 | 0.1301 | 1.9822 | 0.0684 | 0.0705 |
| | Residuals | 27 | 1.7718 | 0.0656 | 0.9316 | | |
| | Total | 28 | 1.9019 | 1.0000 | | | |
| TA | Stage | 1 | 0.2502 | 0.2502 | 1.9250 | 0.1017 | 0.1054 |
| | Residuals | 17 | 2.2092 | 0.1300 | 0.8983 | | |
| | Total | 18 | 2.4593 | 1.0000 | | | |
| **WA** | **Stage** | **1** | **0.5188** | **0.5188** | **6.2490** | **0.2066** | **0.0004** |
| | **Residuals** | **24** | **1.9926** | **0.0830** | **0.7934** | | |
| | **Total** | **25** | **2.5115** | **1.0000** | | | |

**Supplementary Table 7:** PERMANOVA analysis with vegetational variables (Plant cover and Trees) for the four chronosequences experimenting large community changes through soil development. Significant relationships (p<0.05) in bold

| Cronosequence | | Df | SS | MS | Pseudo F | $R^2$ | P-value |
|---|---|---|---|---|---|---|---|
| **CH** | **Plant cover** | **1** | **0.2842** | **0.2842** | **4.9275** | **0.1543** | **0.0015** |
| | **Residuals** | **27** | **1.5575** | **0.0577** | **0.8457** | | |
| | **Total** | **28** | **1.8417** | | | | |
| | **Trees** | **1** | **0.1518** | **0.1518** | **2.4245** | **0.0824** | **0.0283** |
| | **Residuals** | **27** | **1.6899** | **0.0626** | **0.9176** | | |
| | **Total** | **28** | **1.8417** | **1.0000** | | | |
| **ALPS** | Plant cover | 1 | 0.0395 | 0.0395 | 0.3678 | 0.0181 | 0.8534 |
| | Residuals | 20 | 2.1464 | 0.1073 | 0.9819 | | |
| | Total | 21 | 2.1859 | | | | |
| | **Trees** | **1** | **0.6756** | **0.6756** | **8.9474** | **0.3091** | **0.0014** |
| | **Residuals** | **20** | **1.5102** | **0.0755** | **0.6909** | | |
| | **Total** | **21** | **2.1859** | **1.0000** | | | |
| **WA** | Plant cover | 1 | 0.1391 | 0.1391 | 1.4074 | 0.0554 | 0.2045 |
| | Residuals | 24 | 2.3723 | 0.0988 | 0.9446 | | |
| | Total | 25 | 2.5115 | | | | |
| | **Trees** | **1** | **0.6063** | **0.6063** | **7.6369** | **0.2414** | **0.0001** |
| | **Residuals** | **24** | **1.9052** | **0.0794** | **0.7586** | | |
| | **Total** | **25** | **2.5115** | **1.0000** | | | |
| **MI** | **Plant cover** | **1** | **0.5096** | **0.5096** | **4.0226** | **0.0804** | **0.0128** |
| | **Residuals** | **46** | **5.8270** | **0.1267** | **0.9196** | | |
| | **Total** | **47** | **6.3365** | **1.0000** | | | |
| | **Trees** | **1** | **1.1201** | **1.1201** | **9.8771** | **0.1768** | **0.0001** |
| | **Residuals** | **46** | **5.2164** | **0.1134** | **0.8232** | | |
| | **Total** | **47** | **6.3365** | **1.0000** | | | |
